# Structural basis for the therapeutic advantage of dual and triple agonists at the human GIP, GLP-1 or GCG receptors

**DOI:** 10.1101/2021.07.29.454286

**Authors:** Fenghui Zhao, Qingtong Zhou, Zhaotong Cong, Kaini Hang, Xinyu Zou, Chao Zhang, Yan Chen, Antao Dai, Anyi Liang, Qianqian Ming, Mu Wang, Linan Chen, Peiyu Xu, Rulue Chang, Wenbo Feng, Tian Xia, Yan Zhang, Beili Wu, Dehua Yang, Lihua Zhao, H. Eric Xu, Ming-Wei Wang

## Abstract

Glucose homeostasis, regulated by glucose-dependent insulinotropic polypeptide (GIP), glucagon-like peptide-1 (GLP-1) and glucagon (GCG) is critical to human health. Several multi-targeting agonists at GIPR, GLP-1R or GCGR, developed to maximize metabolic benefits with reduced side-effects, are in clinical trials to treat type 2 diabetes and obesity. To elucidate the molecular mechanisms by which tirzepatide, a GIPR/GLP-1R dualagonist, and peptide 20, a GIPR/GLP-1R/GCGR triagonist, manifest their superior efficacies over monoagonist such as semaglutide, we determined cryo-electron microscopy structures of tirzepatide-bound GIPR and GLP-1R as well as peptide 20-bound GIPR, GLP-1R and GCGR The structures reveal both common and unique features for the dual and triple agonism by illustrating key interactions of clinical relevance at the atomic level. Retention of glucagon function is required to achieve such an advantage over GLP-1 monotherapy. Our findings provide valuable insights into the structural basis of functional versatility and therapeutic supremacy of tirzepatide and peptide 20.

## Introduction

Glucose-dependent insulinotropic polypeptide (also known as gastric inhibitory peptide, GIP), glucagon-like peptide-1 (GLP-1) and glucagon (GCG) are peptide hormones responsible for glucose homeostasis^1,2^. Their cognate receptors, GIPR, GLP-1R and GCGR, belong to class B1 G protein-coupled receptor (GPCR) family. Successful application of various GLP-1 mimetics to treat type 2 diabetes mellitus (T2DM) and obesity highlights the clinical value of this group of drug targets^3^. However, development of GIPR- and GCGR-based therapeutics has encountered drawbacks due to the complexity of physiology associated with GIP and GCG^4-6^. For example, GIP stimulates insulin secretion but also increases GCG levels^7,8^, while the latter has a parallel role in elevating energy expenditure and blood glucose^9^.

It was reported that the weight loss property (5-10%) of GLP-1 analogs is hampered by dose-dependent side-effects^10^. Chimeric peptides consisting of amino acids from GIP and GLP-1 were then designed to maximize their metabolic benefits^11^. Additional consideration was given to GCG for its role in energy expenditure^12^. Therefore, multi-targeting or unimolecular peptides possessing combinatorial agonism at GIPR, GLP-1R and GCGR have been extensively explored and more than a dozen peptides including two GIPR/GLP-1R dualagonists, ten GLP-1R/GCGR dualagonists and five GIPR/GLP-1R/GCGR triagonists have entered into clinical development (Fig. S1a, Supplementary Table 1)^13^. Of them, two pioneered unimolecular agonists, tirzepatide (LY3298176) and peptide 20 (MAR423) have attracted significant attention from both academic and industrial communities (Fig. 1a). Tirzepatide is an investigational once-weekly GIPR/GLP-1R dualagonist^14^ with a profound therapeutic superiority in reducing blood glucose and body weight beyond several approved drugs such as semaglutide^15^ and dulaglutide^16^ in multiple head-to-head clinical trials. Peptide 20, a GIPR/GLP-1R/GCGR triagonist (currently in phase 1 clinical trial)^17^ with balanced potency at the three receptors, is evolved from a GLP-1R/GCGR dualagonist^18^ through iterative sequence refinement and modification (Fig. S1b)^14^. It reversed glucose dysregulation without detrimental effects on metabolically healthy animals and reduced body weight, lowered fasting blood glucose, decreased glycosylated hemoglobin (HbA1C), improved glucose tolerance, and protected pancreatic islet architecture in diabetic fatty Zucker rats^14,19,20^.

**Fig. 1.**
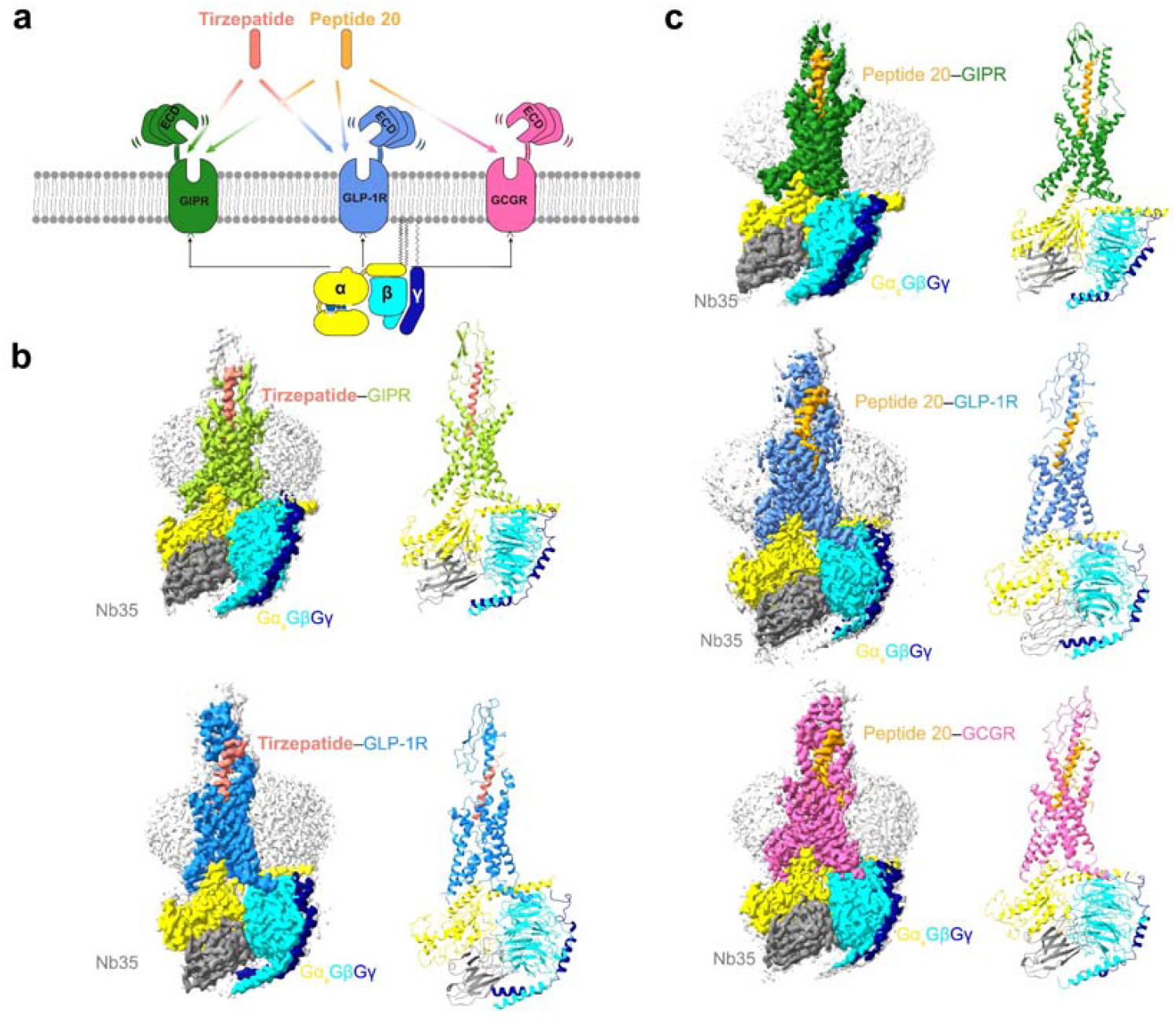
Cryo-EM structures of tirzepatide and peptide 20-bound GIPR, GLP-1R and GCGR in complex with G_s_. **a**, Unimolecular peptides tirzepatide and peptide 20 possess distinct combinatorial agonism at GIPR, GLP-1R and GCGR. **b**, Cryo-EM maps (left) and structural models (right) of tirzepatide-bound GIPR (top) and GLP-1R (bottom) in complex with G_s_. The sharpened cryo-EM density map at the 0.243 threshold shown as light gray surface indicates a micelle diameter of 10 nm. The colored cryo-EM density map is shown at the 0.424 threshold. The tirzepatide is shown in salmon, GIPR in yellow green, GLP-1R in dodger blue, Gα_s_ in yellow, Gβ subunit in cyan, Gγ subunit in navy blue and Nb35 in gray. **c**, Cryo-EM maps (left) and structural models (right) of peptide 20-bound GIPR (top), GLP-1R (middle) and GCGR (bottom) in complex with G_s_. The sharpened cryo-EM density map at the 0.228 threshold shown as light gray surface indicates a micelle diameter of 11 nm. The colored cryo-EM density map is shown at the 0.576 threshold. The peptide 20 is shown in orange, GIPR in forest green, GLP-1R in blue, GCGR in hot pink, Gα_s_ in yellow, Gβ subunit in cyan, Gγ subunit in navy blue and Nb35 in gray.

To understand molecular mechanisms of the dual and triple agonism conferred by tirzepatide and peptide 20, we determined five cryo-electron microscopy (cryo-EM) structures, including GIPR and GLP-1R bound with tirzepatide and GIPR, GLP-1R and GCGR bound with peptide 20, all in complex with G_s_ proteins at global resolutions of 3.4 Å, 3.4 Å, 3.1 Å, 3.0 Å and 3.5 Å, respectively. Integrated with pharmacological and clinical data, this work reveal the structural basis of peptide recognition by each receptor and provide important insights into therapeutic benefits resulted from combinatorial agonism.

## Results

### Overall structure

The tirzepatide–GIPR–G_s_, tirzepatide–GLP-1R–G_s_, peptide 20–GIPR–G_s_, peptide 20–GLP-1R–G_s_ and peptide 20–GCGR–G_s_ structures were determined by the single-particle cryo-EM approach with overall resolutions of 3.4 Å, 3.4 Å, 3.1 Å, 3.0 Å, and 3.5 Å, respectively (Fig. 1b,c, Figs. S2-6, Table S1, Supplementary Figure 1, Supplementary Table 2). Apart from the α-helical domain of Gα_s_, the presence of bound tirzepatide and peptide 20, individual receptor and heterotrimeric G_s_ in respective complex was clearly visible in all five EM maps, thereby allowing unambiguous modeling of the secondary structure and side chain orientation of all major components of the complexes (Fig. S6).

Tirzepatide has two non-coded amino acid residues at positions 2 and 13 (Aib, α-aminoisobutyric acid), and is acylated on K20^P^ (P indicates that the residue belongs to the peptide) with a γGlu-2×OEG linker and C18 fatty diacid moiety. The first 30 and 29 amino acids of tirzepatide were modelled for the tirzepatide–GIPR–G_s_ and tirzepatide–GLP-1R–G_s_ complexes, respectively.

Peptide 20 contains two modifications: A2^P^ with Aib and K10^P^ that is covalently attached by a 16-carbon acyl chain (palmitoyl; 16:0) via a gamma carboxylate (γE spacer)^14^. The γE spacer and palmitic acid (C16:0) were well resolved in the final models of peptide 20–GCGR–G_s_ and peptide 20–GLP-1R–G_s_, while only the γE spacer was modelled for peptide 20–GIPR–G_s_ with high-resolution features. The first 30, 29, and 28 amino acids of peptide 20 were modelled for the peptide 20–GIPR–G_s_, peptide 20–GLP-1R–G_s_ and peptide 20–GCGR–G_s_ complexes, respectively.

As shown in Fig. 2a, the tirzepatide–GIPR–G_s_ and peptide 20–GIPR–G_s_ complex structures closely resembled that of the GIP–GIPR–G_s_ complex^21^ with Cα root mean square deviation (RMSD) values of 0.5 and 0.4 Å, respectively. Notable conformational differences were observed in the positions of peptide C-terminal half and the surrounding ECL1 and ECD, indicative of GIPR-associated ligand specificity. Through two mutations (M14^P^L and H18^P^A), the dense contacts between ECL1 (residues 194 to 211) and GIP were disrupted by peptide 20, as seen from the buried surface area that decreased from 406 Å^2^ for GIP to 278 Å^2^ for peptide 20. Consequently, ECL1 adopted a more relaxed conformation, making peptide 20 straighter by shifting its tip toward the TMD core by 4.2 Å (measured by the Cα of L27^P^). Similar movement was also seen for the C-terminal half of tirzepatide (2.1 Å measured by the Cα of I27^P^). As far as the N terminus is concerned, GIP and tirzepatide were stabilized by massive contacts with TMD core through a common N terminus (Y1^P^-A/Aib2^P^-E3^P^), while that of peptide 20 (H1^P^-Aib2^P^-Q3^P^) formed weaker interactions with TMD core by abolishing the hydrogen bond with Q224^3.37b^ (class B GPCR numbering in superscript)^22^, salt bridge with R183^2.60b^ and hydrophobic contacts with V227^3.40b^ (Fig. 2b). Such deficiency of peptide 20 was rescued by the introduction of T7^P^ (hydrogen bond with R190^2.67b^), lipidated K10^P^ and Y13^P^ that contributed additional contacts with GIPR not observed in GIP^21^. The hydrogen bond between T7^P^ and R190^2.67b^ was also found in the tirzepatide–GIPR–G_s_ complex.

**Fig. 2.**
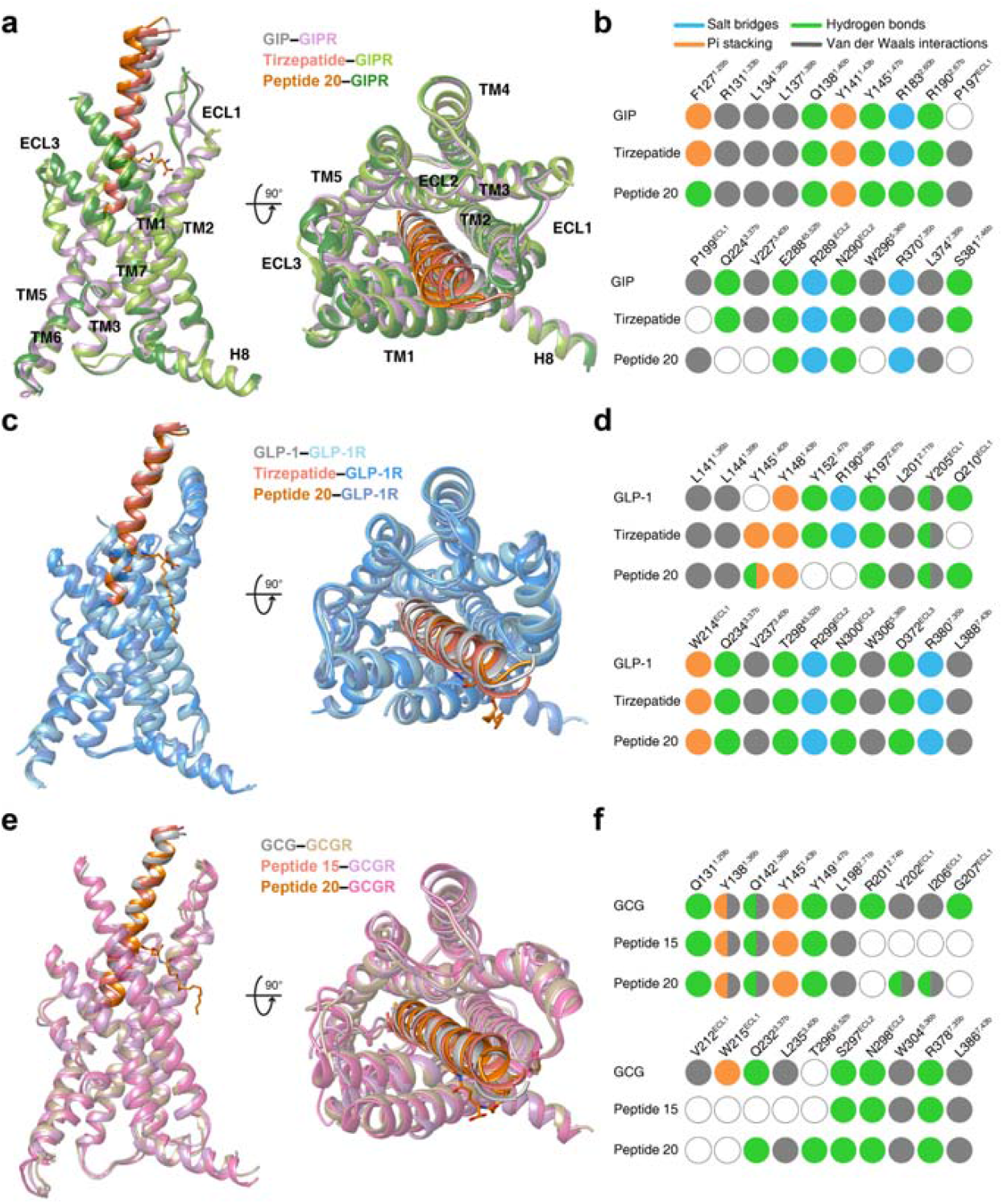
Structural comparison of GIPR, GLP-1R and GCGR bound by mono-, dual and triple agonists. **a**, Structural comparison of GIP–GIPR–G_s_ ^21^, tirzepatide–GIPR–G_s_and peptide 20–GIPR–G_s_. Receptor ECD and G protein are omitted for clarity. **b**, Comparison of residue interactions employed by GIPR to recognize GIP, tirzepatide and peptide 20, described by fingerprint strings encoding different interaction types of the surrounding residues in each peptide. Color codes are listed on the top panel. Residues that show no interaction with ligands are displayed as white circles. **c**, Structural comparison of GLP-1–GLP-1R–G_s_ ^23^, tirzepatide–GLP-1R–G and peptide 20–GLP-1R–G_s_. Receptor ECD and G protein are omitted for clarity. **d**, Comparison of residue interactions that GLP-1R employed to recognize GLP-1, tirzepatide and peptide 20, described by fingerprint strings encoding different interaction types of the surrounding residues in each peptide. **e**, Structural comparison of GCG–GCGR–G_s_ ^4^, peptide 15–GCGR–G_s_ ^24^ and peptide 20–GCGR–G_s_ . Receptor ECD and G protein are omitted for clarity. **f**, Comparison of residue interactions that GCGR employed to recognize GCG, peptide 15 and peptide 20, described by fingerprint strings encoding different interaction types of the surrounding residues in each peptide.

The structures of tirzepatide- and peptide 20-bound GLP-1R are highly similar to that bound by GLP-1^23^, with Cα RMSD of 0.8 Å and 0.7 Å, respectively (Fig. 2c). The bound peptides (GLP-1, tirzepatide and peptide 20) overlapped well and penetrated into the receptor TMD core by an identical angle and orientation, thereby exploiting a similar ligand recognition pattern for most residues except for a few positions that have distinct amino acids (Fig. 2c, Supplementary Tables 3, 4). The substitution (Y10^P^ in tirzepatide) and modification (lipidated K10^P^ in peptide 20) stabilized the binding of dual and triple agonists by newly-formed interactions with residues surrounding the TM1-TM2 cleft, a phenomenon unseen in the case of GLP-1^23^. Meanwhile, some favorable interactions in GLP-1 recognition were absent for both tirzepatide (Y13^P^A decreased the hydrophobic interactions with TM1, E21^P^A broke the hydrogen bond with Q210^ECL1^) and peptide 20 (E3^P^Q eliminated the salt bridge with R190^2.60b^) (Fig. 2d). Interestingly, the residues at multiple positions (12, 16, 17, 20, 21, 24 and 28) of the unimolecular agonists are highly solvent-accessible and of limited contact with GLP-1R, allowing them to employ distinct amino acids from GLP-1 without altering GLP-1R signaling profiles. As a comparison, superimposing either GIP or GCG with GLP-1 analogs suggest that they have potential steric clashes with ECL1 of GLP-1R via H18^P^ of GIP and R18^P^ of GCG. Two residues with shorter side-chains (I7^P^ and A13^P^) in GIP further weakened its binding to GLP-1R, consistent with the distinct cross-reactivity features of GIP and GCG with GLP-1R^5,6^.

Superimposing the structures of GCGR–G_s_ bound by GCG^4^, peptide 15 (GLP-1R and GCGR dual agonist)^24^ and peptide 20 reveals that these three peptides adopt a similar binding pose: a single continuous helix that penetrates into the TMD core through their N-terminal halves (residues 1 to 15), while the C-terminal halves (residues 16 to 30) are recognized by the ECD, ECL1 and TM1 (Fig. 2e). Given that both peptide 15 and peptide 20 are modified forms of GCG (differed by 7 residues), ligand recognition patterns are highly conserved across the three peptides except for a few positions. For example, by choosing alanine at position 18 instead of arginine in GCG, peptide 20 lost the cation-pi stacking with W215^ECL1^ and hydrogen bond with Q204^ECL1^, thereby allowing its outward movement toward ECL1 and leading to the formation of another hydrogen bond (D21^P^-I206^ECL1^) (Fig. 2f). Probably due to the lack of complementary interacting residues, aligning GIP or GLP-1 to GCG significantly loosened the dense compact between GCG and GCGR by removing one hydrogen bond (Y10^P^(GCG)/Y10^P^(GIP)/V16^P^(GLP-1)-Q142^1.40b^(GCGR)) and pi-pi stacking (Y13^P^(GCG)/A13^P^(GIP)/Y19^P^(GLP-1)-Y138^1.36b^(GCGR)) and by repulsing the interaction between Y1^P^(GIP) and I235^3.40b^(GCGR). These observations receive the support of our current and previous functional data showing that both GIP and GLP-1 were unable to activate GCGR (Supplementary Table 5)^5,6^.

Collectively, the binding mode comparison of the three peptides bound by the same receptor demonstrate common structural features in ligand recognition and distinct conformational adaptability of GIPR, GLP-1R and GCGR in response to different agonist stimulation.

### Recognition of tirzepatide

The tirzepatide–GIPR–G_s_ and tirzepatide–GLP-1R–G_s_ exhibit a similar peptide-receptor binding interface, where distinct structural features were observed at ECL1, ECL3 and the extracellular tips of TM1 and TM3 (Fig. 3a). GIPR-bound tirzepatide is rotated by 8.3° compared to that in complex with GLP-1R, such a movement shifted its C terminus toward TMD core by 5.2 Å (measured by the Cα of I27^P^). The N-terminal region of tirzepatide (residues 1 to 10) in GIPR and GLP-1R overlapped well with the formation of a network of extensive interactions with multiple conserved residues (Y^1.43b^, Y^1.47b^, R190/K197^2.67b^, Q^3.37b^, V^3.40b^, N290/N300^ECL2^, R^7.35b^ and I378/L388^7.43b^) (Fig. 3b-e, Supplementary Tables 3, 6). Notably, the inward movement of GIPR R300^5.40b^ contributed one hydrogen bond with T5^P^ (Fig. 3b, f). The middle region of tirzepatide in GLP-1R was stabilized by the peptide-ECD-ECL1-ECL2 interface through both a polar network (T298^45.52^-S11^P^-Y205^ECL1^-R299^ECL1^-D15^P^-L32^ECD^-S31^ECD^-Q19^P^) and a complementary nonpolar network with ECD (L32, V36, W39 and Y88) and ECL1 (W214) via F22^P^, W25^P^, L26^P^ (Fig. 3c). As a comparison, the ECL1 of GIPR partially unwound with the presence of three proline residues (P195^ECL1^, P197^ECL1^ and P199^ECL1^), resulting in reduced interactions between ECL1 and tirzepatide compared to that in GLP-1R (Fig. 3b). However, the α-helical extension in TM1 of GIPR provides additional residues for tirzepatide recognition including one hydrogen bond (Y10^P^ and Q138^1.40b^) and a stacking interaction (K16^P^ and F127^1.29b^). The acylation on K20^P^ by γGlu-2×OEG linker and C18 fatty diacid moiety that enables enhanced binding to plasma albumin and extended the peptide half-life *in viv*o^25^ were not resolved in both structures, indicating a high conformational flexibility, in line with the recently published cryo-EM structure of semaglutide-bound GLP-1R^26^and our molecular dynamics (MD) simulation results (Fig. S7a-c). Consistently, the non-acylated tirzepatide maintained high affinity and potency to both GLP-1R and GIPR as tirzepatide (Fig. S2f, g).

**Fig. 3.**
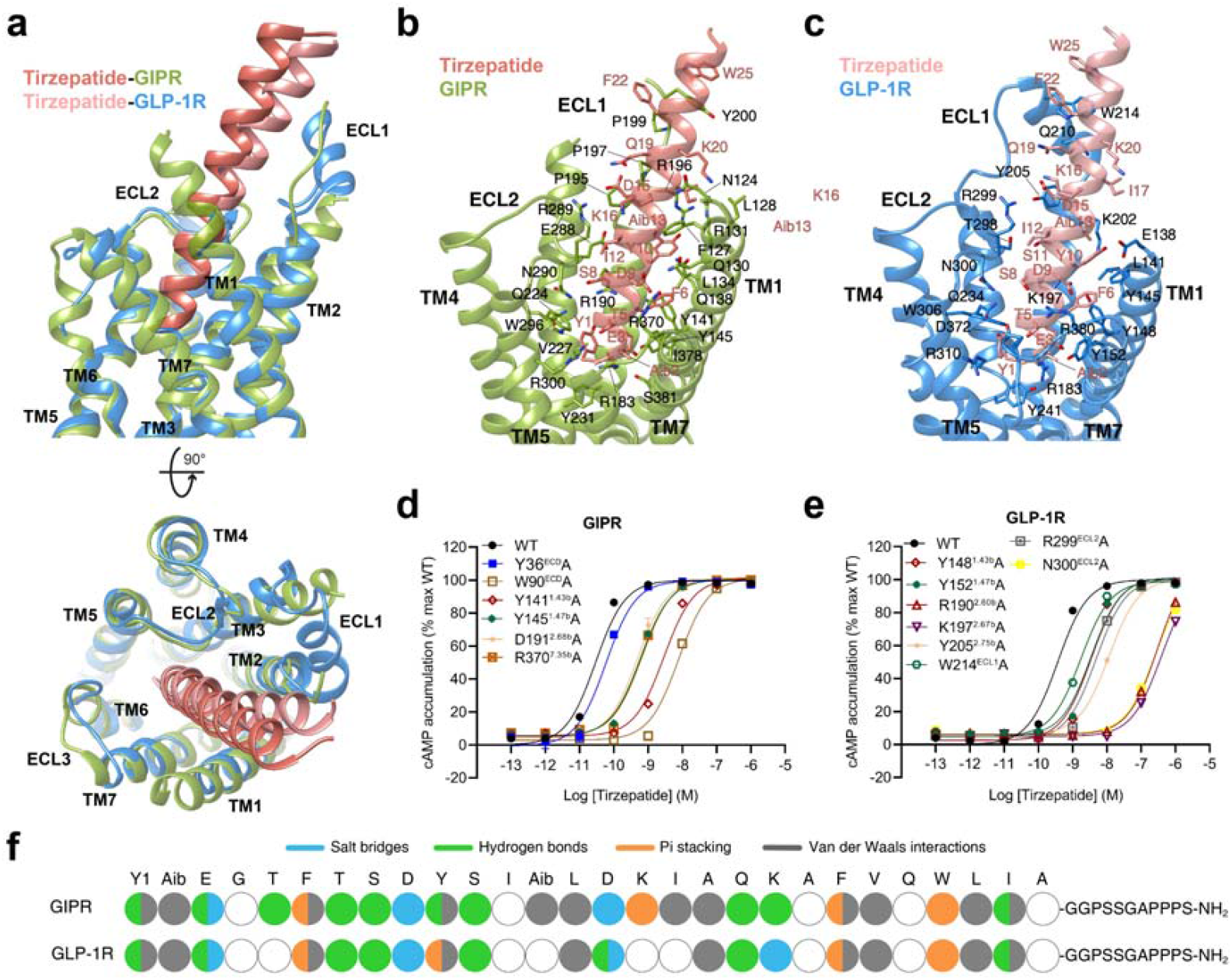
Molecular recognition of tirzepatide by GIPR and GLP-1R. **a**, Structural comparison of tirzepatide–GIPR–G_s_ and tirzepatide–GLP-1R–G_s_. Receptor ECD and G protein are omitted for clarity. **b**, Interactions between tirzepatide (salmon) and the TMD of GIPR (yellow green). Residues involved in interactions are shown as sticks. **c**, Interactions between tirzepatide (light salmon) and the TMD of GLP-1R (dodger blue). Residues involved in interactions are shown as sticks. **d-e**, Effects of receptor mutations on tirzepatide-induced cAMP accumulation. Data shown are means ± S.E.M. of at least three independent experiments performed in quadruplicate. **f**, The peptide recognition modes are described by fingerprint strings encoding different interaction types of the surrounding residues in each receptor. Residues that show no interaction with receptors are displayed as white circles. Color codes are listed on the top panel. WT, wild-type.

### Peptide 20 recognition

Superimposition of the TMDs of GIPR, GLP-1R and GCGR bound by peptide 20 shows that the three receptors employed conserved residues in the lower half of the TMD pocket to recognize the well-overlapped peptide N-terminal region (residues 1 to 11), while the peptide C terminus engaged by ECL1, the N-terminal α-helix of ECD and the extracellular tip of TM1 display receptor-specific positions and orientations (Fig. 4, Fig. S8). Accompanying the inward movement of GIPR ECL1 by 6.4 Å relative to that of GCGR (measured by Cα of G202^ECL1^ in GIPR and G207^ECL1^ in GCGR), the C terminus of peptide 20 bound by GIPR shifted toward TMD core by 8.1 Å (measured by Cα of L27^P^) and consequently pushed the extracellular tip of TM1 moving toward TM7 by 2.8 Å (measure by Cα of the residues at 1.29b). ECL1 and ECD of the three receptors coincidently constructed a complementary binding groove for the entrance of the C terminus of peptide 20 through multiple hydrophobic residues (A19^P^, F22^P^, V23^P^, W25^P^, L26^P^ and L27^P^). However, several additional interactions were observed in GLP-1R (S11^P^-Y205^ECL1^ and D21^P^-Q210^ECL1^) and GCGR (D15^P^-Y202^ECL1^ and D21^P^-I206^ECL1^), but not in GIPR (Fig. 4b-h, Supplementary Tables 4, 7, 8).

**Fig. 4.**
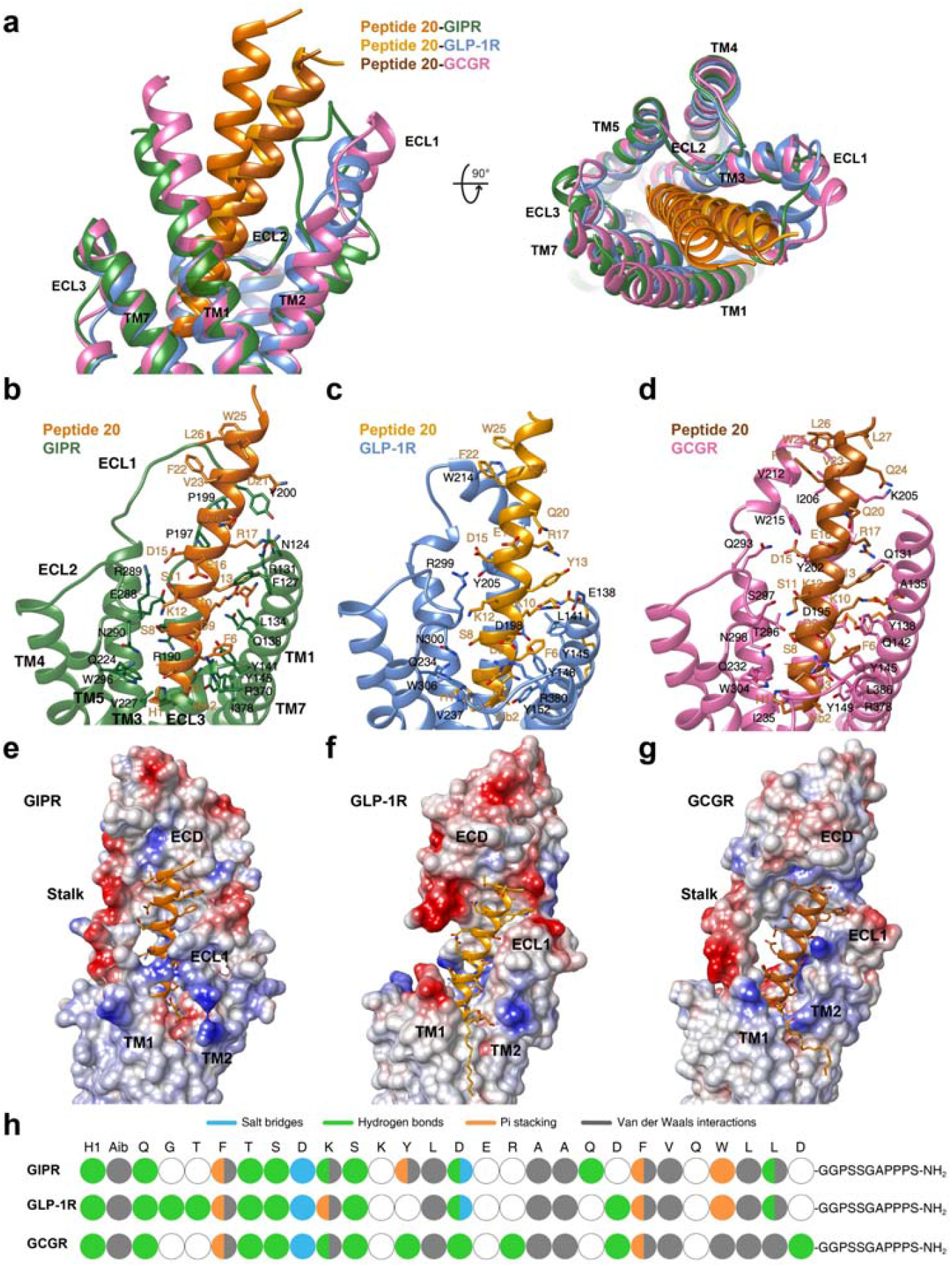
Molecular recognition of peptide 20 by GIPR, GLP-1R and GCGR. **a**, Structural comparison of peptide 20–GIPR–G_s_, peptide 20–GLP-1R–G_s_ and peptide 20–GCGR–G_s_. Receptor ECD and G protein are omitted for clarity. **b-d**, Interactions between peptide 20 and the TMDs of GIPR (forest green), GLP-1R (blue), and GCGR (hot pink). Residues involved in interactions are shown as sticks. **e-g**, Electrostatic surface representations of the receptor for each of the peptide-receptor complex, with the peptides shown as ribbon and sticks. Electrostatic surface potential was calculated in Chimera according to Coulomb’s law and contoured at ± 10 kT e^−1^. Negatively and positively charged surface areas are colored red and blue, respectively. **h**, The peptide recognition modes are described by fingerprint strings encoding different interaction types of the surrounding residues in each receptor. Color codes are listed on the top panel. Residues that show no interaction with receptors are displayed as white circles.

Notably, strong cryo-EM densities were observed in the crevices between TM1 and TM2 of the three complexes (Fig. 5a-c). They were connected to the side-chain end of K10^P^ of peptide 20, allowing unambiguous assignment of the binding sites of lipidated K10^P^ with a 16-carbon palmitic acid through a γ-carboxylate spacer (Fig. 5d-f). Such a modification on K10^P^ greatly stabilized the peptide binding through extensive contacts with both receptors and lipid membrane. For GCGR, the lipidated K10^P^ contributed three hydrogen bonds (with S139^1.37b^, Q142^1.40b^ and R199^2.72b^), extensive hydrophobic contacts (with V143^1.41b^, T146^1.44b^, L192^2.65b^ and V193^2.66b^) and lipid membrane where the 16-carbon palmitic chain implanted (Fig. 5d-f). Removal of these contacts by GCGR triple mutant (Q142A+D195A+R199A) markedly reduced peptide 20 potency by 93-fold (Fig. 5g). For GLP-1R, the γ-carboxylate spacer formed two hydrogen bonds (with Y145^1.40b^ and D198^2.68b^), and the 16-carbon palmitic chain terminus dropped down along TM1 with the formation of massive hydrophobic interactions with I146^1.41b^, T149^1.44b^, V150^1.45b^, A153^1.48b^ and L154^1.49b^. Similar phenomenon was also observed in GIPR. Consistently, our MD simulations found that the γ-carboxylate spacer stably inserted into the TM1-TM2 cleft and the 16-carbon palmitic chain is deeply buried in the receptor-lipid interface, contributing massive contacts to stabilize the complexes (Fig. S7d, e). The importance of K10^P^ lipidation receives the support of our structure-activity relationship study where peptide 20 without K10^P^ lipidation reduced the receptor-mediated cAMP accumulation by 8,709-fold and 660-fold for GIPR and GCGR, respectively, but inappreciably influenced that of GLP-1R (Fig. 5h). These results suggest that specific modification of peptide is equally significant to sequence optimization in term of demonstration of a desired polypharmacology of a unimolecular dual or triple agonist.

**Fig. 5.**
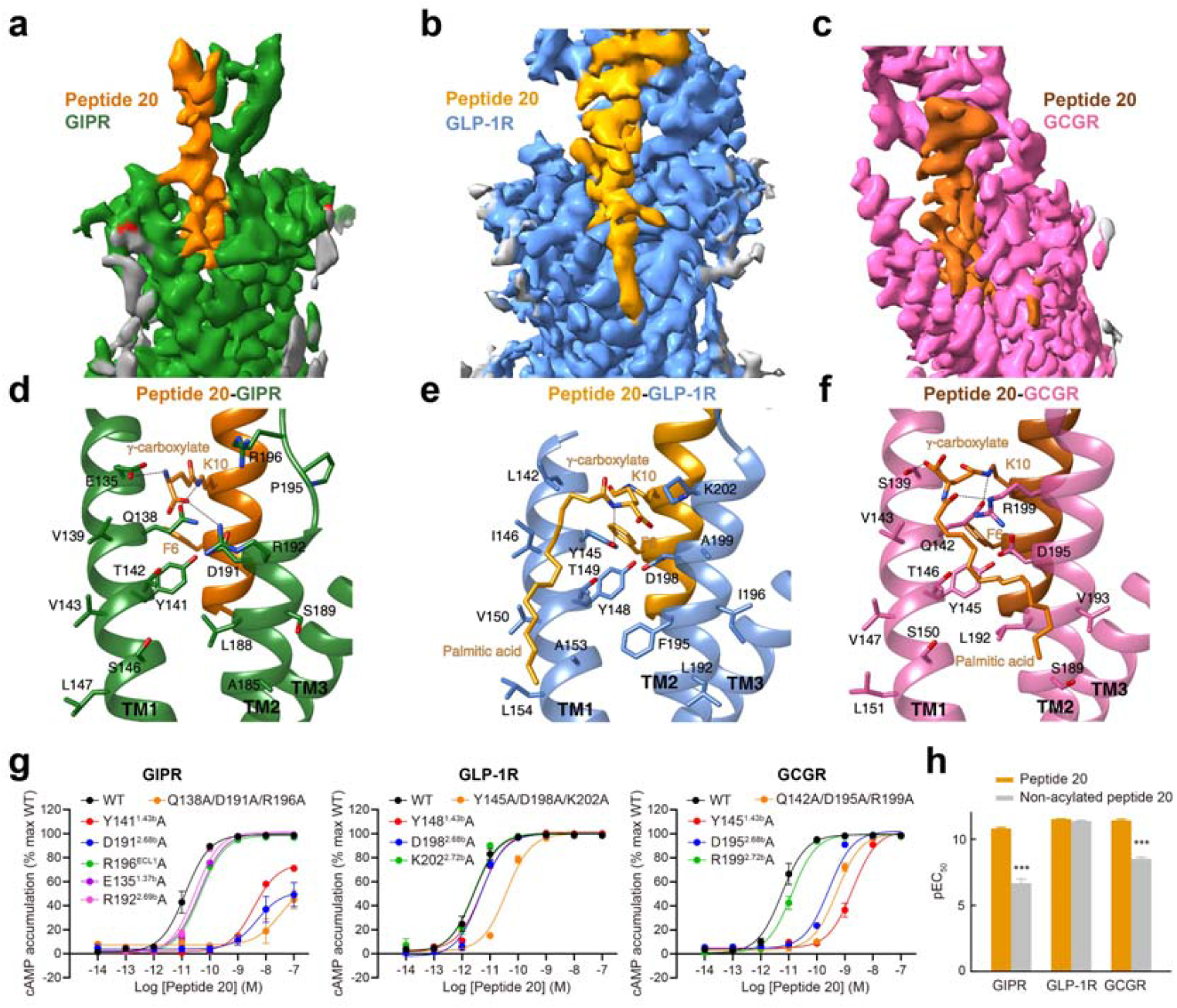
Structural and functional feature of lipidated K10^P^ of peptide 20. **a-c**, Close-up views of the crevices between TM1 and TM2 displayed by cryo-EM maps of peptide 20-bound GIPR (**a**), GLP-1R (**b**) and GCGR (**c**). Continuous electron densities connected to K10 in peptide 20 were observed in the three peptide 20-bound receptor–G_s_ complexes. **d-f**, Interactions between lipidated K10^P^ and the TM1-TM2 crevice of GIPR (**d**), GLP-1R (**e**) and GCGR (**f**), with interacting residues shown in sticks. Hydrogen bonds are shown with dashed lines. **g**, Effects of receptor mutations on peptide 20-induced cAMP accumulation. Data shown are means ± S.E.M. of at least three independent experiments performed in quadruplicate. **h**, Effects of K10 lipidation on peptide 20-induced cAMP accumulation. The bar graph represents the average pEC_50_ (that is, −logEC_50_) measured from three independent experiments performed in quadruplicate. Statistically significant differences were determined with a two-tailed Student’s t test. ***P< 0.001. WT, wild-type.

### Receptor activation

Despite the existence of unique structural features among the ligand-binding pockets of GIPR, GLP-1R and GCGR, both tirzepatide and peptide 20 triggered receptor conformational changes similar to that induced by GLP-1 or GCG^4,23^ and distinct from the inactive or *apo* GLP-1R and GCGR structures (Fig. S9)^27,28^. Compared to the inactive GCGR, the extracellular tip of TM7 in peptide 20-bound GCGR moved outward by 5.1 Å (measured by Cα atom of L377^7.34b^) and the α-helical structure of the extracellular half of TM6 was partially unwounded. In the intracellular side, a sharp kink located in the conserved Pro^6.47b^-X-X-Gly^6.50b^ motif pivoted the intracellular tip of TM6 to move outwards by 19.3 Å (measured by Cα atom of K344^6.35b^), slightly higher than that seen with the GCG–GCGR–G_s_ (17.7 Å)^4^. This, in conjunction with the movement of TM5 towards TM6, opened up the cytoplasmic face of GCGR to accommodate G protein coupling. Similar conformational change was also observed in the tirzepatide–GIPR–G_s_, tirzepatide–GLP-1R–G_s_, peptide 20–GIPR–G_s_ and peptide 20–GLP-1R–G_s_ complexes, compared to peptide-free *apo* GLP-1R structure^27^. At the residue level, signaling initiation by either peptide 20, tirzepatide or endogenous peptide hormones rendered a common arrangement of residue contacts for the three receptors^29,30^, including the reorganization of the central polar network that located just below the peptide binding site, opening of the hydrophobic packing to favor the formation of TM6 kink at the PXXG motif and the rearrangement of two polar networks (HETX motif and TM2-6-7-helix 8) at the cytoplasmic face.

### G protein coupling

Comparison of the two tirzepatide- and three peptide 20-bound GPCR–G_s_ complex structures with that of other class B1 GPCR family members reveals a high similarity in the G protein binding interface, suggesting a common mechanism for G_s_ engagement^4,29,31-34^ (Fig. 6a). These complexes are anchored by the α5 helix of Gα_s_, which fits to the cytoplasmic cavity formed by TMs 2, 3, 5, 6, 7 and intracellular loop 1 (ICL1). Besides, H8 contributes several polar interactions with the Gβ subunit. There are some receptor- and ligand-specific structural features displayed by ICL2. For peptide 20-bound GCGR, its ICL2 moved downward and made extensive polar and nonpolar contacts with the binding groove formed by the αN helix, β1 strand and α5 helix of Gα_s_, resulting in an ICL2–Gα_s_ interface area of 799 Å^2^, significantly larger than that of GLP-1R (396 Å^2^) or GIPR (416 Å^2^) (Fig. 6b). Different from the dipped down side-chain conformation observed in GLP-1-bound GLP-1R^23^, F257^3.60b^ in the peptide 20–GLP-1R–G_s_complex rotated its side-chain upwards (Fig. 6c). Furthermore, E262^ICL2^ was reoriented ∼90° from an outside facing position to a position pointing to Gα_s_, thus introducing a hydrogen bond with Q35^GαHN^ (Fig. 6d). Similar G protein interface was also observed in the tirzepatide-bound GLP-1R except for the orientation of E262^ICL2^ that is closer to that of GLP-1. In the case of peptide 20- and tirzepatide-bound GIPR complexes, the side-chain of E253^ICL2^contributed one salt bridge with K34^GαHN^, not observed in the peptide 20-bound GLP-1R and GCGR complexes (Fig. 6e).

**Fig. 6.**
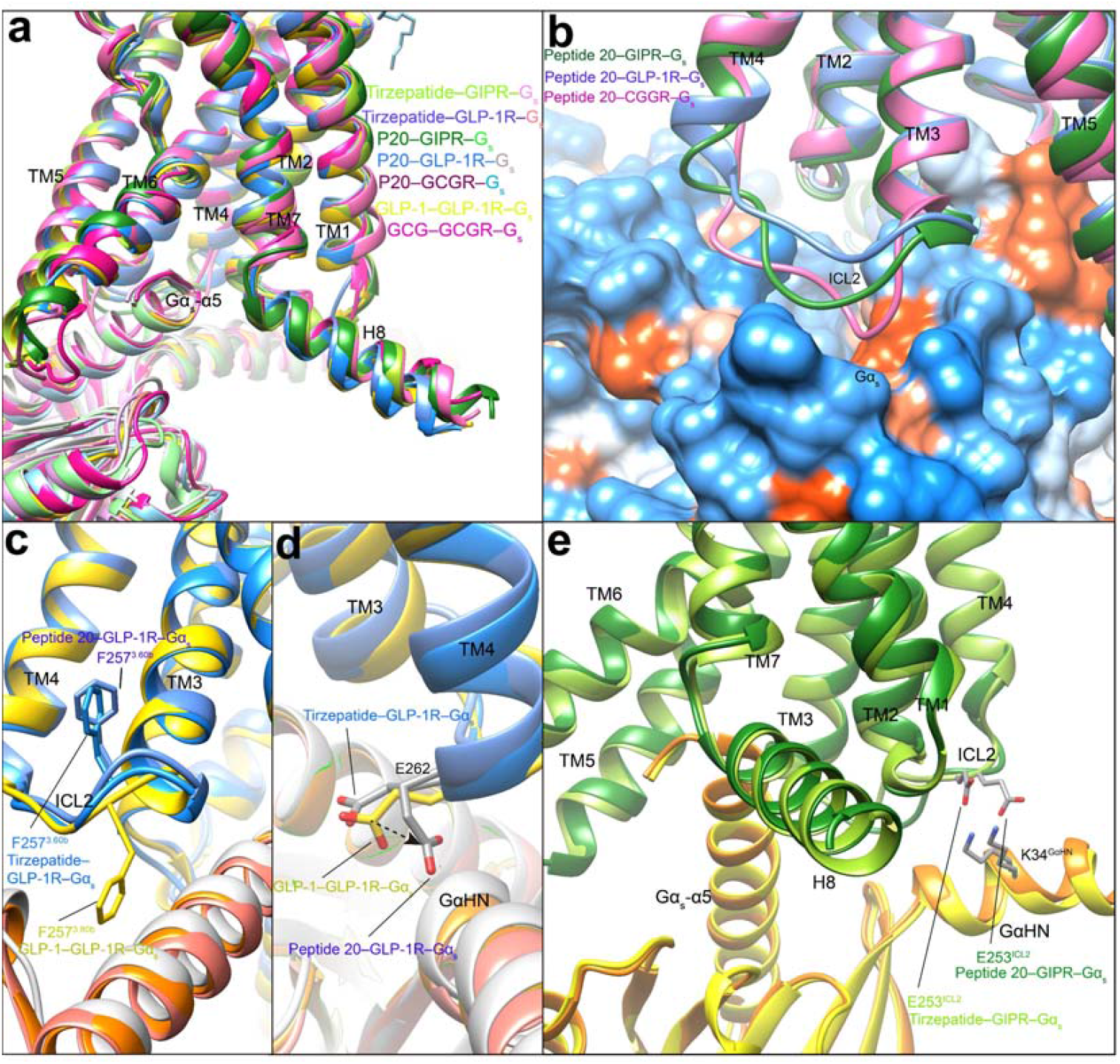
G protein coupling of unimolecular agonist-bound GIPR, GLP-1R and GCGR. **a**, Comparison of G protein coupling among GIPR, GLP-1R and GCGR^4,21,23^. The Gα_s_ α5-helix of the Gα_s_ Ras-like domain inserts into an intracellular crevice of receptor’s TMD. The receptors and G proteins are colored as the labels. **b**, Comparison of ICL2 conformation in the peptide 20-bound GIPR, GCGR and GLP-1R. **c**, Comparison of F257^3.60b^ conformation in the GLP-1R bound by GLP-1, tirzepatide and peptide 20. **d**, Comparison of E262^ICL2^ conformation in the GLP-1R bound by GLP-1, tirzepatide and peptide 20. **e**, Comparison of E253^ICL2^ conformation in the GIPR bound by tirzepatide and peptide 20. Residues involved in interactions are shown as sticks. Polar interactions are shown as black dashed lines.

### Efficacy superiority

The superior therapeutic efficacy of tirzepatide over approved selective GLP-1 analogs were reported recently^16,35^, whereas the outcome of clinical trials on peptide 20 is not available in the literature. The five high-resolution cryo-EM structures reported here, together with abundant structural and pharmacological data of monospecific peptides documented previously^4,21,23,26,36^, provide us an excellent opportunity to analyze the molecular basis of the superior clinical efficacy presented by unimolecular agonists.

Semaglutide and tirzepatide share two common substitutions (Aib8^P^ and acylated K26^P^ by C18 diacids via a γGlu-2×OEG linker, numbered according to GLP-1 and semaglutide whose first N-terminal residues are at position 7 while that of tirzepatide is at position 1) introduced to reduce degradation by dipeptidyl peptidase-4 (DPP-4) and to prolong their half-lives by enhanced binding to plasma albumin (Fig. 7a)^37^. Besides, there is only one residue in semaglutide (R34^P^) that is different from GLP-1 but does neither form any interaction with GLP-1R^26^ nor affect receptor binding and signaling^25^. However, tirzepatide has 14 unique amino acids (engineered from the GIP sequence) and an amidated exenatide-like C terminus as opposed to GLP-1 which allow the peptide to possess a GIPR binding ability equivalent to GIP(1–42) and to steadily interact with GLP-1R with a reduced potency compared to GLP-1^23^ (Fig. 2a-d). Like GLP-1, semaglutide is not able to bind or activate GIPR. These findings were confirmed by GIPR or GLP-1R mediated cAMP accumulation assays (Fig. 7b-c)^35^. Of note is that tirzepatide was reported to cause biased signaling at GLP-1R in favor of cAMP response over β-arrestin recruitment^35^. The combined activation of GIPR and GLP-1R by tirzepatide not only improved both glucose-dependent insulin secretion and glucose tolerance in mice^38^, but also showed significantly better efficacy than semaglutide and dulaglutide with regard to glucose control and weight loss^15,16^.

**Fig. 7.**
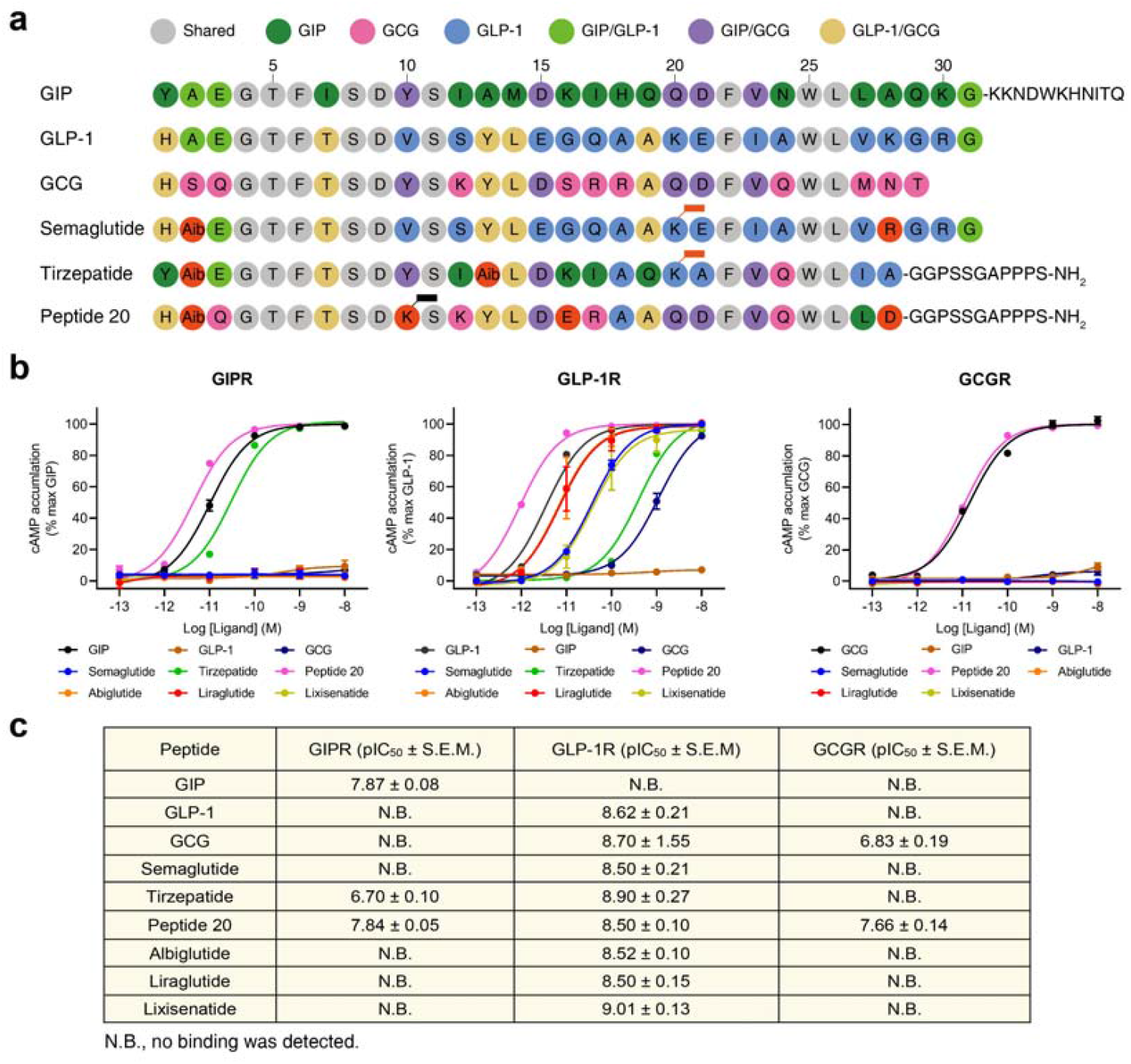
Structure-basis of receptor selectivity demonstrated by tirzepatide, peptide 20 and GLP-1 analogs. **a**, Amino acid sequences of endogenous agonists, unimolecular agonists and approved GLP-1 analogs including semaglutide. Residues are colored according to sequence conservation among GIP, GLP-1 and GCG. Aib, aminoisobutyric acid. Semaglutide and tirzepatide are conjugated by a C20 fatty diacid moiety via a linker connected to the lysine residue at position 20, while peptide 20 is covalently attached by a 16-carbon acyl chain (palmitoyl; 16:0) via a γ-carboxylate spacer at K10^P^. **b**, Receptor signaling profiles of endogenous agonists, unimolecular agonists and approved drug GLP-1 analogs including semaglutide. Data shown are means ± S.E.M. of at least three independent experiments performed in quadruplicate. **c**, Receptor binding profiles of endogenous agonists, unimolecular agonists and approved GLP-1 analogs. Data shown are means ± S.E.M.

It is known that peptide 20 potently reversed metabolic disorders in rodent models of obesity and diabetes, characteristic of increased energy expenditure and elevated circulating FGF21 levels as a result of GCGR agonism^14,19^. Peptide 20 utilizes a N terminus (the first 11 residues) that is highly conserved across GIP, GLP-1 and GCG to interact with the lower half of the TMD pocket of the three receptors consisting of conserved residues such as L/Y^1.36b^ (hydrophobic with K10^P^), Q/Y^1.40b^ (hydrogen bond with K10^P^), Y^1.43b^ (stacking with F6^P^), Y^1.47b^(hydrogen bond with Q3^P^), Q^3.37b^ (hydrogen bond with H1^P^), ECL2 (hydrogen bond with S8^P^), R^7.35b^ (salt bridge with D9^P^), I/L^7.43b^ (hydrophobic with Aib2^P^) and L^7.43b^ (hydrophobic with F6^P^) (Figs. 2, 4b-d, 7a). A similar approach was applied to the design of peptide 20’s C terminus that occupies the hydrophobic binding groove of ECD, with residues (A19^P^, F22^P^, V23^P^, W25^P^, L26^P^ and L27^P^) adopted from GIP, GLP-1 and GCG (Figs. 4e-g, 7a)^39,40^. To accommodate the upper half of the TMD pocket formed by ECL1 and the extracellular tips of TM1 and TM2 that diversified in both sequence and conformation across the three receptors, peptide 20 employs distinct “barcodes” (patterns of amino acids) to recognize specific region of a given receptor (Fig. 4h). For GIPR whose ECL1 was loosely compacted by peptide 20, three residues (Y13^P^-L14^P^-D15^P^) strengthened the peptide-binding interface by forming a hydrogen bond with F127^1.29b^ and a salt bridge with R289^ECL2^, significantly stronger than that observed in GLP-1R and GCGR. Alternatively, another three residues (D21^P^-F22^P^-W25^P^) compacted well with the ordered ECL1 of GLP-1R via a hydrogen bond with Q210^ECL1^ and packing with W214^ECL1^. Two hydrogen bonds (D15^P^-Y202^2.75b^ and R17^P^-Y202^2.75b^) were only seen in GCGR.

The most impressive structural feature of peptide 20 is the lipidated K10^P^ by a 16-carbon palmitic acid through a γ-carboxylate spacer, which perfectly inserted into TM1-TM2 crevice and made extensive contacts with both receptors and lipid membrane to stabilize the binding poses (Fig. 5). These observations disclose a combined mechanism that uses conserved residues for ligand recognition and specific “barcodes” to accommodate conformations unique to each receptor, leading to a highly potent and balanced unimolecular triple agonist for GIPR, GLP-1R and GCGR^14^ with a cAMP signaling potency similar to that of GIP, GLP-1 and GCG (Fig. 7b).

## Discussion

Due to the central roles exerted by the three metabolically related peptide hormone receptors (GIPR, GLP-1R and GCGR) in the management of T2DM and obesity, the concept of combinatorial agonism or polypharmacology to synergize metabolic actions and maximize therapeutic benefits has been explored in the past decade with remarkable preclinical and clinical achievements. The 3-dimensional structures of GCGR, GLP-1R and GIPR solved previously helped us better understand the molecular basis of ligand recognition and receptor activation of these important class B1 GPCRs^21,28,41-43^. In this paper, we report five cryo-EM structures of two well-recognized unimolecular agonists (tirzepatide and peptide 20) in complex with individual receptors and G_s_ proteins. The structural basis of their superior clinical efficacies relative to monospecific agonists such as semaglutide is elucidated. Our results provide an atomic level visualization of the molecular action of unimolecular agonists on three cognate receptors and offer valuable information for the design of better drugs to combat metabolic disease.

Superimpositions of the two tirzepatide- and three peptide 20-bound structures to the three receptors bound by the endogenous ligands (GIP, GLP-1 and GCG) showed that the five peptides all adopt a single continuous helix, with the well-overlapped N terminus penetrating to the TMD core stabilized by conserved interactions, while the C terminus anchors the ECD, ECL1 and ECL2 in a receptor- and ligand-specific manner. With the presence of three proline residues (P195^ECL1^, P197^ECL1^ and P199^ECL1^), the ECL1 of GIPR presents a notable conformational adaptability in recognition of different agonists, a phenomenon that was not seen with that of GLP-1R and GCGR as their binding pockets exhibit less flexibility when recognizing the peptides through a combination of common segment that contributes to conserved interactions and distinct sequences that govern receptor selectivity. The distinct sequences that tirzepatide and peptide 20 employed, respectively, to recognize GIPR or GLP-1R are obviously different: the former was primarily based on the GIP sequence with engineered GLP-1 activity^38^, whereas the latter was derived from a GLP-1R/GCGR dualagonist in conjunction with GIP agonism^14^. Such a sequence and receptor binding divergence may consequently alter pharmacological and clinical outcomes. Clearly, distinct sequence and structural features of tirzepatide and peptide 20 allow them to exert combinatorial agonism at two or more receptors at the same time thereby maximize the benefit of polypharmacology and minimize the limitation of mono-targeting.

Both GIP and GLP-1 are released upon nutrient ingestion to promote insulin secretion by pancreatic β-cells. However, they have opposed effects on circulating GCG levels^7,15^. GIPR activation also has different roles in lipid metabolism from that of GLP-1^44^. Maintenance of GCG action might be a key to the superior therapeutic efficacy of tirzepatide^15,16,45^. Structurally, the binding of tirzepatide to GIPR reshaped the ECL1 conformation relative to that of GIP, but made no change in the GLP-1R structure. As far as peptide 20 is concerned, the peptide binding pocket of both GLP-1R and GCGR closely resembled that of GLP-1 and GCG bound structures, where notable conformational change was only observed in the ECL1 of GIPR. These differences in structural plasticity or rigidity among the three receptors give clues to further optimize unimolecular agonists using complementary amino acids to target common regions of individual receptors and distinct sequences to confer receptor selectivity.

Unlike tirzepatide that retains GCG function via counteracting with that of GLP-1 through activation of GIPR, peptide 20 is capable of activating GCGR directly. Consistent with the effects of GCGR in increasing lipolysis and thermogenesis besides elevating blood glucose levels, preclinical studies have found that peptide 20 improved energy metabolism and hepatic lipid handling without exacerbating preexisting hyperglycemia^14^. Peptide 20 was developed through a series of optimizing processes based on GCGR agonism in diet-induced obese mice, concluding that the ideal metabolic benefits of triagonism predominantly depend on fine-tuning the GCG component^14^. The structures reveal that lipidation at K10 of peptide 20 allows the hydrophobic acyl tail to interact with the TMD region of all three receptors, providing a new clue for peptidic ligand design. From the perspective of precision medicines, combinatorial agonism might be precisely designed to reflect pharmacological profiles of individual receptors such that diabetic patients at different disease stages could be prescribed with different unimolecular agonists to take personalized therapeutic advantages.

## Supporting information

Seven PDBs and their maps

## Methods

### Cell lines

*Spodoptera frugiperda* 9 (*Sf*9) (Invitrogen) and High Five™ insect cells (Expression Systems) were cultured in ESF 921 serum-free medium (Expression Systems) at 27°C and 120 rpm. Human embryonic kidney 293 cells containing SV40 large T-antigen (HEK293T) were cultured in DMEM (Gibco) supplemented with 10% (v/v) fetal bovine serum (FBS, Gibco), 1 mM sodium pyruvate (Gibco) and 100 units/mL penicillin and 100 μg/mL streptomycin at 37°C in 5% CO_2_. Chinese hamster ovary (CHO-K1) cells were cultured in F-12 (Gibco) containing 10% FBS, 100 units/mL penicillin and 100 μg/mL streptomycin at 37°C in 5% CO_2_. For cAMP and receptor expression assays, HEK293T cells were seeded into 6-well cell culture plates at a density of 7 × 10^5^ cells per well. For whole-cell binding assay, CHO-K1 cells were seeded into 96-well fibronectin-treated cell culture plates at a density of 3 × 10^4^ cells per well. After overnight incubation, cells were transfected with GIPR, GLP-1R or GCGR construct using Lipofectamine 2000 transfection reagent (Invitrogen). Following 24 h culturing, the transfected cells were ready for use.

### Construct

The human GIPR DNA (Genewiz) with one mutation (T345F) was cloned into the pFastBac vector (Invitrogen) with its native signal peptide replaced by the haemagglutinin (HA) signal peptide. A BRIL fusion protein was added at the N-terminal of the ECD with a TEV protease site and 2GSA linker between them. C-terminal 45 amino acids (Q422-C466) of the receptor were truncated. LgBiT was added at the end of helix 8 with a 15-amino acid (15AA) polypeptide linker in between, followed by a TEV protease cleavage site and an OMBP-MBP tag. A dominant-negative bovine Gα_s_ (DNGα_s_) construct with 9 mutations (S54N, G226A, E268A, N271K, K274D, R280K, T284D, I285T and A366S)^58,59^ was used to help stabilize the tirzepatide–GIPR–G_s_ complex. Meanwhile, a DNGα_s_ construct with 8 mutations (S54N, G226A, E268A, N271K, K274D, R280K, T284D and I285T) was used to help stabilize the peptide 20–GIPR–G_s_ complex^34,59^. Rat Gβ1 was cloned with a C-terminal SmBiT34 (peptide 86 or HiBiT, Promega) connected with a 15AA polypeptide linker. The modified rat Gβ1 and bovine Gγ2 were both cloned into a pFastBac vector. The construct and various mutants of human GIPR were cloned into pcDNA3.1 vector for cAMP accumulation and whole-cell binding assays.

The human GLP-1R was modified with its native signal sequence (M1-P23) replaced by the HA signal peptide to facilitate receptor expression. To obtain a GLP-1R–G_s_ complex with good homogeneity and stability, we used the NanoBiT tethering strategy, in which the C terminus of GLP-1R was directly attached to LgBiT subunit followed by a TEV protease cleavage site and a double MBP tag. Rat Gβ1 was the same as the construct used in the GIPR structure determination. The Gα_s_ (DNGα_s_ with 9 mutations) used to stabilize the tirzepatide–GLP-1R–G_s_ complex was the same as that employed for the tirzepatide–GIPR–G_s_ complex. A dominant-negative human Gα_s_ (DNGα_s_) with 8 mutations (S54N, G226A, E268A, N271K, K274D, R280K, T284D and I285T) was generated as previously described to limit G protein dissociation^59^. The constructs were cloned into both pcDNA3.1 and pFastBac vectors for functional assays in mammalian cells and protein expression in insect cells, respectively. Other constructs including the full-length and various mutants of human GLP-1R were cloned into pcDNA3.1 vector for cAMP accumulation and whole-cell binding assays.

The human GCGR gene was cloned into pFastBac1 vector with GP64 promoter at the N terminus to enhance the protein yield. Forty-five residues (H433-F477) were truncated at the C terminus to improve the thermostability and an affinity tag, HPC4 tag, was added to the C terminus (GP64-HA-GCGR-GSGS linker-HPC4). Gα_s_ (DNGα_s_ with 8 mutations) was modified as above to stabilize the interaction with βγ subunits. The rat Gβ1 and bovine Gγ2 were used in the structure determination.

Additionally, we used an engineered G_s_ (mini-G_s_) protein to stabilize the non-acylated tirzepatide (the side-chain was removed at C20) bound GIPR or GLP-1R as described previously^60^.

### Protein expression

Baculoviruses containing the above complex constructs were prepared by the Bac-to-Bac system (Invitrogen). For the tirzepatide–GIPR–G_s_ and non-acylated tirzepatide–GIPR–mini-G_s_ complexes, GIPR and DNGα_s_ or mini-G_s_ heterotrimer were co-expressed in High Five™ cells. Briefly, insect cells were grown in ESF 921 culture medium (Expression Systems) to a density of 3.2 × 10^6^ cells/mL. The cells were then infected with BRIL-TEV-2GSA-GIPR(22-421)T345F-15AA-LgBiT-TEV-OMBP-MBP, DNGα_s_ or mini-G_s_, Gβ1-peptide 86 and Gγ2, respectively, at a ratio of 1:4:4:4. For the peptide 20–GIPR–G_s_ complex, GIPR and G_s_ heterotrimer were co-expressed in High Five™ cells grown in ESF 921 culture medium (Expression Systems) to a density of 3.2 × 10^6^ cells/mL. The cells were then infected with BRIL-TEV-2GSA-GIPR(22-421)T345F-15AA-LgBiT-TEV-OMBP-MBP, DNGα_s_, Gβ1-peptide 86 and Gγ2, respectively, at a ratio of 1:3:3:3. After 48 h incubation at 27°C, the cells were collected by centrifugation and stored at -80°C until use.

The GLP-1R-LgBiT-2MBP, DNGα_s_ or mini-G_s_, Gβ1-peptide 86 and Gγ2 were co-expressed at multiplicity of infection (MOI) ratio of 1:1:1:1 by infecting *Sf*9 cells at a density of 3.0 × 10^6^ cells/mL. Other operations are the same as GIPR.

The GCGR construct, DNGα_s_ and Gβ1 and Gγ2 were co-expressed in High Five™ cells and infected with four separate baculoviruses at a ratio of 4:1:1:1. Other operations are the same as GIPR.

### Nb35 expression and purification

Nanobody-35 (Nb35) with a 6× his tag at the C-terminal was expressed in the periplasm of *E. coli* BL21 (DE3) cells. Briefly, Nb35 target gene was transformed in the bacterium and amplified in TB culture medium with 100 μg/mL ampicillin, 2 mM MgCl_2_, 0.1 % (w/v) glucose at 37°C, 180 rpm. When OD600 reached 0.7-1.2, 1 mM IPTG was added to induce expression followed by overnight incubation at 28°C. The cell pellet was then collected under 4°C and stored at -80°C. Nb35 was purified by size-exclusion chromatography (SEC) using a HiLoad 16/600 Superdex 75 column (GE Healthcare) with running buffer containing 20 mM HEPES, 100 mM NaCl, pH 7.4. Fractions of Nb35 were concentrated to ∼2 mg/mL and quickly frozen in the liquid nitrogen with 10% glycerol and stored in -80°C.

### Complex formation and purification

For the tirzepatide–GIPR–G_s_ complex, cell pellets were lysed in a buffer containing 20 mM HEPES, 100 mM NaCl, pH 7.4, 10 mM MgCl_2_, 1 mM MnCl_2_ and 10% glycerol supplemented with protease inhibitor cocktail, EDTA-free (TragetMol). Cell membranes were then collected by ultracentrifugation at 4°C, 90,000 *g* for 35 min. A buffer consisting of 20 mM HEPES, 100 mM NaCl, pH 7.4, 10 mM MgCl_2_, 1 mM MnCl_2_ and 10% glycerol was used to re-suspend the collected membranes. To assemble the GIPR–G_s_ complex, 15 μM tirzepatide (GL Biochem) was added to the preparation accompanied by 100 μM TCEP, 25 mU/mL apyrase (Sigma-Aldrich), 20 μg/mL Nb35 and 100 U salt active nuclease (Sigma-Aldrich) supplemented with protease inhibitor cocktail for 1.5 h incubation at room temperature (RT). The membrane was then solubilized with 0.5% (w/v) lauryl maltose neopentylglycol (LMNG, Anatrace) and 0.1% (w/v) cholesterol hemisuccinate (CHS, Anatrace) with additional 2 μM tirzepatide for 3 h at 4°C. The supernatant was isolated by centrifugation at 90,000 *g* for 35 min and the solubilized complex was incubated with amylose resin (NEB) for 2.5 h at 4°C. The resin was collected by centrifugation at 550 *g* and loaded onto a gravity flow column. The resin in the column was first washed with 5 column volumes (CVs) of buffer containing 20 mM HEPES, pH 7.4, 100 mM NaCl, 10% (v/v) glycerol, 5 mM MgCl_2_, 1 mM MnCl_2_, 25 μM TCEP, 5 μM tirzepatide, 0.1% (w/v) LMNG and 0.02% (w/v) CHS. After this, the resin was further washed with 25 CVs of buffer containing 20 mM HEPES, pH 7.4, 100 mM NaCl, 10% (v/v) glycerol, 5 mM MgCl_2_, 1 mM MnCl_2_, 25 μM TCEP, 5 μM tirzepatide, 0.03% (w/v) LMNG, 0.01% (w/v) glyco-diosgenin (GDN, Anatrace) and 0.008% (w/v) CHS. The protein was then incubated with a buffer consisting of 20 mM HEPES, pH 7.4, 100 mM NaCl, 10% (v/v) glycerol, 5 mM MgCl_2_, 1 mM MnCl_2_, 25 μM TCEP, 50 μM tirzepatide, 20 μg/mL Nb35, 0.03% (w/v) LMNG, 0.01% (w/v) GDN, 0.008% (w/v) CHS and 30 μg/mL His-tagged TEV protease on the column overnight at 4°C. The flow through was collected and concentrated to 500 μL using a 100 kDa filter (Merck Millipore). SEC was performed by loading the protein onto Superose 6 Increase 10/300GL (GE Healthcare) column with running buffer containing 20 mM HEPES, pH 7.4, 100 mM NaCl, 10 mM MgCl_2_, 100 μM TCEP, 5 μM tirzepatide, 0.00075% (w/v) LMNG, 0.00025% (w/v) GDN, 0.0002% (w/v) CHS and 0.00025% digitonin (Anatrace). The tirzepatide–GIPR–G_s_ complexes were collected and concentrated for cryo-EM analysis.

For the non-acylated tirzepatide–GIPR–mini-G_s_ complex, the operations of the purification were the same as the tirzepatide–GIPR–G_s_ complex, except that the peptide was replaced by the non-acylated tirzepatide. The complex samples were concentrated to 14-16 mg/mL for cryo-EM analysis.

For the tirzepatide–GLP-1R–G_s_ complex, cells were suspended in 20 mM HEPES, pH 7.4, 100 mM NaCl and 10% (v/v) glycerol in the presence of protease inhibitor cocktail. Complex was formed by adding 10 mM MgCl_2_, 1 mM MnCl_2_, 50 mU/mL apyrase, 30 μM tirzepatide, 100 μM TCEP and 10 μg/mL Nb35 to the cell lysate and incubated at RT for 1.5 h. Cell membranes were solubilized by adding 0.5% (w/v) LMNG supplemented with 0.1% (w/v) CHS at 4°C for 2 h, followed by centrifugation at 65,000 *g* for 30 min at 4°C. The supernatant was taken to bind with amylose resin for 2 h at 4°C. After packing, the column was washed with buffer containing 20 mM HEPES, pH 7.4, 100 mM NaCl, 10% (v/v) glycerol, 5 μM tirzepatide, 25 μM TCEP, 5 mM MgCl_2_, 1 mM MnCl_2_, 0.1% (w/v) LMNG and 0.02% (w/v) CHS first (10 CVs), and then with decreased concentrations of detergents, 0.03% (w/v) LMNG, 0.01% (w/v) GDN and 0.006% (w/v) CHS (20 CVs). TEV enzyme was added to the resin and kept at 4°C overnight to remove the OMBP-MBP tag. The complex was eluted from the resin and concentrated to 500 μL using a 100 kDa MWCO Amicon Ultra Centrifugal Filter. SEC was carried out by loading the protein sample to Superdex 200 Increase 10/300GL (GE Healthcare) to obtain the monomer complex. The column was pre-equilibrated with 20 mM HEPES, pH 7.4, 100 mM NaCl, 5 μM tirzepatide, 100 μM TCEP, 2 mM MgCl_2_, 0.00075% (w/v) LMNG, 0.00025% (w/v) GDN, 0.00015% (w/v) CHS and 0.00025% digitonin.

For the non-acylated tirzepatide–GLP-1R–mini-G_s_ complex, the operations of the purification were the same as the peptide 20–GLP-1R–G_s_ complex, except that the peptide was replaced by the non-acylated tirzepatide, and the detergent of SEC running buffer was changed to 0.01% digitonin. The complex samples were concentrated to 16-18 mg/mL for cryo-EM analysis.

For the peptide 20–GIPR–G_s_ complex, the operations of the purification was the same as the tirzepatide–GIPR–G_s_ complex, except that the peptide was replaced by the peptide 20. The complex samples were concentrated to 5-6 mg/mL for cryo-EM analysis.

For the peptide 20–GLP-1R–G_s_ complex, cell pellets were thawed and lysed in a buffer containing 20 mM HEPES, pH 7.5, 100 mM NaCl, 10% (v/v) glycerol, 10 mM MgCl_2_, 1 mM MnCl_2_ and 100 μM TCEP supplemented with EDTA-free protease inhibitor cocktail by dounce homogenization. The complex formation was initiated by the addition of 20 μM peptide 20, 10 μg/mL Nb35 and 25 mU/mL apyrase. After 1.5 h incubation at RT, the membrane was solubilized in the buffer above supplemented with 0.5% (w/v) LMNG and 0.1% (w/v) CHS for 2 h at 4°C. The supernatant was isolated by centrifugation at 65,000 *g* for 30 min and incubated with amylose resin for 2 h at 4°C. The resin was then collected by centrifugation at 500 *g* for 10 min and washed in gravity flow column with 5 CVs of buffer containing 20 mM HEPES, pH 7.5, 100 mM NaCl, 10% (v/v) glycerol, 5 mM MgCl_2_, 1 mM MnCl_2_, 25 μM TCEP, 0.1% (w/v) LMNG, 0.02% (w/v) CHS and 5 μM peptide 20, followed by washing with 15 CVs of buffer containing 20 mM HEPES, pH 7.5, 100 mM NaCl, 10% (v/v) glycerol, 5 mM MgCl_2_, 1 mM MnCl_2_, 25 μM TCEP, 0.03% (w/v) LMNG, 0.01% (w/v) GDN, 0.008% (w/v) CHS and 5 μM peptide 20. The protein was then incubated overnight with TEV protease on the column to remove the C-terminal 2MBP-tag in the buffer above at 4°C. The flow through was collected next day and concentrated with a 100 kDa molecular weight cut-off concentrator. The concentrated product was loaded onto a Superdex 200 increase 10/300 GL column with SEC running buffer containing 20 mM HEPES, pH 7.5, 100 mM NaCl, 10 mM MgCl_2_, 100 μM TCEP, 2 μM peptide 20, 0.00075% LMNG, 0.00025% GDN and 0.0002% (w/v) CHS. The fractions for monomeric complex were collected and concentrated to 15-20 mg/mL for cryo-EM examination.

For the peptide 20–GCGR–G_s_ complex, cell pellets were resuspended in 20 mM HEPES, pH 7.4, 50 mM NaCl, 2 mM MgCl_2_ with protease inhibitor cocktail, EDTA-free, 5 µM peptide 20, 10 μg/mL Nb35 and 25 mU/mL apyrase. The suspension was incubated at RT for 2 h to promote the formation of complexes. Membranes were collected by centrifugation (30,000 rpm) at 4°C for 30 min, and solubilized in 0.5% (w/v) LMNG, 0.1% (w/v) CHS, 10 µM peptide 20, 2 mM MgCl_2_, 100 U salt active nuclease and 25 mU/ml apyrase for 2.5 h at 4°C. Supernatant was collected by centrifugation at 30,000 rpm for 30 min. The GCGR complex was incubated overnight with anti-HPC4 affinity resin in the presence of 2 mM CaCl_2_, washed with 20 CVs of 20 mM HEPES, pH 7.4, 100 mM NaCl, 2 mM MgCl_2_, 2 mM CaCl_2_, 5 μM peptide 20, 0.02% (w/v) LMNG and 0.004% (w/v) CHS, and eluted with 5 CVs of buffer by adding 6 mM EDTA and 5 µM peptide 20. The complexes were concentrated by a molecular weight cut-off concentrator and separated by SEC on a Superose 6 Increase 10/300GL with running buffer containing 20 mM HEPES, pH 7.4, 100 mM NaCl, 2 mM MgCl_2_, 0.01% (w/v) LMNG, 0.002% (w/v) CHS and 5 μM peptide 20. The complex samples were concentrated to 12-14 mg/mL for cryo-EM analysis.

### Structure determination

To prepare high-quality human GIPR–G_s_ complexes, the receptor’s C terminal forty-five amino acids (Q422-C466) were truncated, and the NanoBiT tethering strategy was applied^21,33,34,61^. To enhance the receptor’s expression, a BRIL fusion protein and an optimized maltose binding protein-maltose binding protein tag (OMBP-MBP)^62^ were added to the N and C termini of the receptor to facilitate the receptor stability and expression (Fig. S2a). To solve the tirzepatide–GIPR–G_s_ complex structure, we introduced one mutation (T345F) to stabilize complex assembly (Fig. S3a). This mutation did not affect ligand binding and signaling properties as verified by both cAMP accumulation and receptor binding assays (Fig. S2d).

The tirzepatide–GLP-1R–G_s_ complex was prepared using the same NanoBiT technique to achieve good homogeneity and stability as described previously^43^ (Fig. S2b). Large-scale purification was performed and the complexes were collected by SEC for cryo-EM studies, with all components of the complex identified in SDS-PAGE of the SEC peak (Fig. S3b). Activation of the modified GIPR and GLP-1R constructs by tirzepatide were confirmed by cAMP accumulation and receptor binding assays, showing similar responses to those of the wild-type (WT) receptors (Fig. S3e-h). Acylated and non-acylated tirzepatide displayed reduced potencies in eliciting GIPR- or GLP-1R-mediated cAMP responses (Fig. S2f, g).

Identical GIPR and GLP-1R constructs were used for the complex structure with peptide 20. Large-scale purification was conducted and the peptide 20–GIPR/GLP-1R–G_s_ complexes were collected by SEC for cryo-EM studies, with all components of the complex identified in SDS-PAGE of the SEC peak (Fig. S4a, b). Activation of the modified GIPR and GLP-1R constructs by peptide 20 were confirmed by cAMP accumulation assays, showing similar responses to those of the WT (Fig. S4d, e). To obtain the peptide 20–GCGR–G_s_ complexes, 45 residues (H433-F477) were truncated at the C terminus of the receptor followed by a HPC4 tag^24^ (Fig. S2c). We used a dominant negative form of Gα_s_ ^30,59^ and nanobody 35 (Nb35) that binds across the Gα:Gβ interface^63^ to enhance protein stability. Purified complex was resolved as a monodisperse peak on SEC, with all components of the complex identified in SDS-PAGE of the SEC peak (Fig. S4c). The modified GCGR construct had a lower potency than that of the WT but did not significantly affect the binding affinity and cAMP signaling of GCG (Fig. S4f).

### Data acquisition and image processing

The purified tirzepatide–GIPR–G_s_–Nb35 complex at a concentration of 18-20 mg/mL was mixed with 100 μM tirzepatide at 4°C and applied to glow-discharged holey carbon grids (Quantifoil R1.2/1.3, Au 300 mesh) that were subsequently vitrified by plunging into liquid ethane using a Vitrobot Mark IV (ThermoFisher Scientific). A Titan Krios equipped with a Gatan K3 Summit direct electron detector was used to acquire cryo-EM images. The microscope was operated at 300 kV accelerating voltage, at a nominal magnification of 46,685× in counting mode, corresponding to a pixel size of 1.071 Å. Totally, 5,434 movies were obtained with a defocus range of -1.2 to -2.2 μm. An accumulated dose of 80 electrons per Å^2^ was fractionated into a movie stack of 36 frames.

The purified tirzepatide–GLP-1R–G_s_–Nb35 complex (3 μL at about 20 mg/mL) was applied to a glow-discharged holey carbon grid (Quantifoil R1.2/1.3) and blotted subsequently. Sample-coated grids were vitrified by plunging into liquid ethane using a Vitrobot Mark IV (ThermoFisher Scientific). Automatic data collection was performed on a Titan Krios equipped with a Gatan K3 Summit direct electron detector. The microscope was operated at 300 kV accelerating voltage, at a nominal magnification of 46,685× in counting mode, corresponding to a pixel size of 1.071 Å. A total of 9,309 movies were obtained with a defocus ranging from -1.2 to -2.2 μm. An accumulated dose of 80 electrons per Å^2^ was fractionated into a movie stack of 45 frames.

The purified peptide 20–GIPR–G_s_–Nb35 complex at a concentration of 5-6 mg/mL was mixed with 100 μM peptide 20 at 4°C and applied to glow-discharged holey carbon grids (Quantifoil R1.2/1.3, Au 300 mesh) that were subsequently vitrified by plunging into liquid ethane using a Vitrobot Mark IV (ThermoFisher Scientific). A Titan Krios equipped with a Gatan K3 Summit direct electron detector was used to acquire cryo-EM images. The microscope was operated at 300 kV accelerating voltage, at a nominal magnification of 46,685× in counting mode, corresponding to a pixel size of 1.071Å. Totally, 3,948 movies were obtained with a defocus range of -1.2 to -2.2 μm. An accumulated dose of 80 electrons per Å^2^ was fractionated into a movie stack of 36 frames.

The purified peptide 20–GCGR–G_s_–Nb35 complex at a concentration of 12-14 mg/mL was mixed with 100 μM peptide 20 at 4°C and applied to glow-discharged holey carbon grids (Quantifoil R1.2/1.3, Au 300 mesh) that were subsequently vitrified by plunging into liquid ethane using a Vitrobot Mark IV (ThermoFisher Scientific). A Titan Krios equipped with a Gatan K3 Summit direct electron detector was used to acquire cryo-EM images. The microscope was operated at 300 kV accelerating voltage, at a nominal magnification of 46,685× in counting mode, corresponding to a pixel size of 1.071Å. Totally, 4,620 movies were obtained with a defocus range of -1.2 to -2.2 μm. An accumulated dose of 80 electrons per Å^2^ was fractionated into a movie stack of 36 frames.

The purified peptide 20–GLP-1R–G_s_–Nb35 complex (3.5 μL) was applied to glow-discharged holey carbon grids (Quantifoil R1.2/1.3, 300 mesh), and subsequently vitrified using a Vitrobot Mark IV (ThermoFisher Scientific) set at 100% humidity and 4°C. Cryo-EM images were collected on a Titan Krios microscope (FEI) equipped with Gatan energy filter and K3 direct electron detector. The microscope was operated at 300 kV accelerating voltage and a calibrated magnification of 46,685× in counting mode, corresponding to a pixel size of 1.071 Å. The total exposure time was set to 7.2 s with intermediate frames recorded every 0.2 s, resulting in an accumulated dose of 80 electrons per Å^2^ with a defocus range of -1.2 to -2.2 μm. Totally, 4,778 images were collected and used for data processing.

The purified non-acylated tirzepatide–GIPR–mini-G_s_–Nb35 complex at a concentration of 14-16 mg/mL was mixed with 100 μM non-acylated tirzepatide at 4°C and applied to glow-discharged holey carbon grids (Quantifoil R1.2/1.3, Au 300 mesh) that were subsequently vitrified by plunging into liquid ethane using a Vitrobot Mark IV (ThermoFisher Scientific). A Titan Krios equipped with a Gatan K3 Summit direct electron detector was used to acquire cryo-EM images. The microscope was operated at 300 kV accelerating voltage, at a nominal magnification of 46,685× in counting mode, corresponding to a pixel size of 1.071Å. Totally, 8,159 movies were obtained with a defocus range of -1.2 to -2.2 μm. An accumulated dose of 80 electrons per Å^2^ was fractionated into a movie stack of 36 frames.

The purified non-acylated tirzepatide–GLP-1R–mini-G_s_–Nb35 complex (3.5 μL) was applied to glow-discharged holey carbon grids (Quantifoil R1.2/1.3, 300 mesh), and subsequently vitrified using a Vitrobot Mark IV (ThermoFisher Scientific) set at 100% humidity and 4°C. Cryo-EM images were collected on a Titan Krios microscope (FEI) equipped with Gatan energy filter and K3 direct electron detector. The microscope was operated at 300 kV accelerating voltage and a calibrated magnification of 46,685× in counting mode, corresponding to a pixel size of 1.071 Å. The total exposure time was set to 7.2 s with intermediate frames recorded every 0.2 s, resulting in an accumulated dose of 80 electrons per Å^2^ with a defocus range of -1.2 to -2.2 μm. Totally, 4,778 images were collected and used for data processing.

Dose-fractionated image stacks were subjected to beam-induced motion correction using MotionCor2.1^64^. A sum of all frames, filtered according to the exposure dose, in each image stack was used for further processing. Contrast transfer function parameters for each micrograph were determined by Gctf v1.06^47^. Automated particle selection and data processing were performed using RELION-3.0 beta2^48^.

For the dataset of the tirzepatide–GIPR–G_s_–Nb35 complex, automated particle selection yielded 4,260,187 particles, which were subjected to reference-free 2D classification, producing 1,771,599 particles with well-defined averages. This subset of particle projections was subjected to a round of 3D classification resulting in one well-defined subset with 870,227 projections. Further 3D classification focusing the alignment on the whole complex produced one high-quality subset accounting for 511,557 particles. These particles were subsequently subjected to CTF refinement and Bayesian polishing, which generated a map with an indicated global resolution of 3.4 Å.

For the dataset of the tirzepatide–GLP-1R–G_s_–Nb35 complex, automated particle selection yielded 4,213,140 particles, which were subjected to reference-free 2D classification, producing 668,880 particles with well-defined averages. This subset of particle projections was subjected to a round of 3D classification resulting in one well-defined subset with 296,989 projections. Further 3D classification focusing the alignment on the whole complex produced one high-quality subset accounting for 125,391 particles. These particles were subsequently subjected to CTF refinement and Bayesian polishing, which generated a map with an indicated global resolution of 3.4 Å.

For the dataset of the peptide 20–GIPR–G_s_–Nb35 complex, automated particle selection yielded 5,322,921 particles. The particles were extracted on a binned dataset with a pixel size of 2.142 Å and were subjected to reference-free 2D classification, producing 4,334,371 particles with well-defined averages. This subset of particle projections was subjected to a round of 3D classification resulting in one well-defined subset with 1,876,783 projections. Further 3D classifications focusing the alignment on the whole complex and the receptor produced one high-quality subset accounting for 255,256 particles. These particles were subsequently subjected to CTF refinement and Bayesian polishing, which generated a map with an indicated global resolution of 3.4 Å.

For the dataset of the peptide 20–GLP-1R–G_s_–Nb35 complex, automated particle selection yielded 4,124,536 particles, which were subjected to reference-free 2D classification, producing 2,354,838 particles with well-defined averages. This subset of particle projections was subjected to a round of 3D classification resulting in one well-defined subset with 1,523,580 projections. Further 3D classifications focusing the alignment on the whole complex and the receptor produced one high-quality subset accounting for 241,786 particles. These particles were subsequently subjected to CTF refinement and Bayesian polishing, which generated a map with an indicated global resolution of 3.1 Å.

For the dataset of the peptide 20–GCGR–G_s_–Nb35 complex, automated particle selection yielded 3,931,945 particles, which were subjected to reference-free 2D classification, producing 917,065 particles with well-defined averages. This subset of particle projections was subjected to a round of 3D classification resulting in one well-defined subset with 578,668 projections. Further 3D classification focusing the alignment on the whole complex produced one high-quality subset accounting for 383,657 particles. These particles were subsequently subjected to CTF refinement and Bayesian polishing, which generated a map with an indicated global resolution of 3.5 Å.

For the dataset of the non-acylated tirzepatide–GIPR–mini-G_s_–Nb35 complex, automated particle selection yielded 7,204,521 particles, which were subjected to reference-free 2D classification, producing 2,718,249 particles with well-defined averages. This subset of particle projections was subjected to a round of 3D classification resulting in one well-defined subset with 2,102,580 projections. Further 3D classification focusing the alignment on the whole complex produced one high-quality subset accounting for 1,251,553 particles. These particles were subsequently subjected to CTF refinement and Bayesian polishing, which generated a map with an indicated global resolution of 3.2 Å.

For the dataset of the non-acylated tirzepatide–GLP-1R–mini-G_s_–Nb35 complex, automated particle selection yielded 5,985,110 particles, which were subjected to reference-free 2D classification, producing 1,723,671 particles with well-defined averages. This subset of particle projections was subjected to a round of 3D classification resulting in one well-defined subset with 906,824 projections. Further 3D classification focusing the alignment on the whole complex produced one high-quality subset accounting for 452,921 particles. These particles were subsequently subjected to CTF refinement and Bayesian polishing, which generated a map with an indicated global resolution of 3.0 Å.

### Model building and refinement

The models of the tirzepatide–GIPR–G_s_ complex and peptide 20–GIPR–G_s_ complex were built using the cryo-EM structure of the GIP–GIPR–G_s_ complex (PDB code: 7DTY)^21^ as the starting point. The models of the tirzepatide–GLP-1R–G_s_ complex and peptide 20–GLP-1R–G_s_ complex were built using the cryo-EM structure of the GLP-1–GLP-1R–G_s_ complex (PDB code: 6×18)^23^ as the starting point. The model of the peptide 20–GCGR–G_s_ complex was built using the cryo-EM structure of the GCG–GCGR–G_s_ complex (PDB code: 6LMK)^4^ as the starting point. The models were docked into the EM density maps using Chimera^51^, followed by iterative manual adjustment and rebuilding in COOT^49^. Real space refinement was performed using Phenix^50^. The model statistics were validated with MolProbity^65^. The final refinement statistics are provided in Table S1.

### cAMP accumulation assay

For GIPR, GLP-1R and GCGR, unimolecular agonist stimulated cAMP accumulation was measured by a LANCE Ultra cAMP kit (PerkinElmer). After 24 h culture, the transfected cells were seeded into 384-well microtiter plates at a density of 3,000 cells per well in HBSS supplemented with 5 mM HEPES, 0.1% (w/v) bovine serum albumin (BSA) and 0.5 mM 3-isobutyl-1-methylxanthine. The cells were stimulated with different concentrations of tirzepatide or peptide 20 for 40 min at RT. Eu-cAMP tracer and ULight™-anti-cAMP were then diluted by cAMP detection buffer and added to the plates separately to terminate the reaction. Plates were incubated at RT for 1 h and the fluorescence intensity measured at 620 nm and 650 nm by an EnVision multilabel plate reader (PerkinElmer).

### Whole-cell binding assay

For GIPR, CHO-K1 cells were cultured in F-12 medium with 10% FBS and seeded at a density of 30,000 cells/well in Isoplate-96 plates (PerkinElmer). The wild-type (WT) or mutant GIPR was transiently transfected using Lipofectamine 2000 transfection reagent as previous described^21^. For homogeneous binding, cells were incubated in binding buffer with a constant concentration of ^125^I-GIP (30 pM, PerkinElmer) and increasing concentrations of unlabeled tirzepatide or peptide 20 (3.57 pM to 1 μM) at RT for 3 h. Following incubation, cells were washed three times with ice-cold PBS and lysed by addition of 50 μL lysis buffer (PBS supplemented with 20 mM Tris-HCl, 1% Triton X-100, pH 7.4). Fifty µL of scintillation cocktail (OptiPhase SuperMix, PerkinElmer) were added and the plates were subsequently counted for radioactivity (counts per minute, CPM) in a MicroBeta^2^ microplate counter (PerkinElmer).

For GLP-1R and GCGR, CHO-K1 cells (3 × 10^4^ per well) were seeded into Isoplate-96 plates and incubated for 24 h at 37°C in 5% CO_2_. They were then washed twice using F-12 with 0.1% BSA, 33 mM HEPES, and incubated for 2 h at 37°C. The medium was removed and ^125^I-GLP-1(7-36)NH_2_ (60 pM) or ^125^I-GCG (40 pM) (PerkinElmer) and increasing concentrations unlabeled tirzepatide or peptide 20 were added for overnight incubation at 4°C. Cells were washed three times with ice-cold PBS and lysed in PBS with 1% Triton X-100, 20 mM Tris-HCl. After addition of scintillation cocktail (PerkinElmer), radioactivity (CPM) was counted on a MicroBeta^2^ microplate counter (PerkinElmer). Data were normalized to the WT response and analyzed using three-parameter logistic equation.

### Receptor expression

Cell surface expression of GIPR, GLP-1R and GCGR were determined by flow cytometry 24 h post-transfection in HEK293T cells. Briefly, approximately 2 × 10^5^ cells were blocked with PBS containing 5% BSA (w/v) at RT for 15 min. After that, cells expressing GIPR and GLP-1R were incubated with 1:300 anti-Flag primary antibody (diluted with PBS containing 5% BSA, Sigma), and those expressing GCGR were incubated with 1:50 anti-GCGR antibody (diluted with PBS containing 5% BSA, Abcam) at RT for 1 h. The cells were then washed three times with PBS containing 1% BSA (w/v) followed by 1 h incubation with 1:1,000 anti-mouse Alexa Fluor 488 conjugated secondary antibody (diluted with PBS containing 5% BSA, Invitrogen) at RT in the dark. After washing three times, cells were resuspended in 200 μL PBS containing 1% BSA for detection by NovoCyte (Agilent) utilizing laser excitation and emission wavelengths of 488 nm and 530 nm, respectively. For each sample, 20,000 cellular events were collected, and the total fluorescence intensity of positive expression cell population was calculated. Data were normalized to the WT receptor.

### Molecular dynamics simulation

Molecular dynamics (MD) simulation was performed by Gromacs 2020.1^52^. The peptide-receptor-complexes were prepared by the Protein Preparation Wizard (Schrodinger 2017-4) with G protein and Nb35 nanobody removed. The receptors were capped with acetyl and methylamide, and the titratable residues were left in their dominant state at pH 7.0. The complexes were embedded in a bilayer composed of 195∼200 POPC lipids and solvated with 0.15 M NaCl in explicitly TIP3P waters using CHARMM-GUI Membrane Builder v3.2.2^54^. The CHARMM36-CAMP force filed^55^ was adopted for protein, peptides, lipids and salt ions. The 16-carbon acyl chain (palmitoyl; 16:0) covalently attached to the side-chain amine of Lys10 in peptide 20 through a γ-carboxylate spacer and the γGlu-2×OEG linker, and C18 fatty diacid moiety that was acylated on Lys26 in tirzepatide were modelled with the CHARMM CGenFF small-molecule force field, program version 1.0.0. The Particle Mesh Ewald (PME) method was used to treat all electrostatic interactions beyond a cut-off of 10 Å and the bonds involving hydrogen atoms were constrained using LINCS algorithm^56^. The complex system was first relaxed using the steepest descent energy minimization, followed by slow heating of the system to 310 K with restraints. The restraints were reduced gradually over 50 ns. Finally, restrain-free production run was carried out for each simulation, with a time step of 2 fs in the NPT ensemble at 310 K and 1 bar using the Nose-Hoover thermostat and the semi-isotropic Parrinello-Rahman barostat^57^, respectively. The buried interface areas were calculated with FreeSASA^53^ using the Sharke-Rupley algorithm with a probe radius of 1.2 Å.

### Statistical analysis

All functional data were presented as means ± standard error of the mean (S.E.M.). Statistical analysis was performed using GraphPad Prism 8 (GraphPad Software). Concentration-response curves were evaluated with a three-parameter logistic equation. The significance was determined with either two-tailed Student’s *t*-test or one-way ANOVA. Significant difference is accepted at P < 0.05.

## Data availability

The atomic coordinates and the electron microscopy maps have been deposited in the Protein Data Bank (PDB) under accession codes: xxx and Electron Microscopy Data Bank (EMDB) accession codes: xxx, respectively. All relevant data are available from the authors and/or included in the manuscript or supplemental data.

## Acknowledgements

We are grateful to Drs. Fan Wu and Fulai Zhou for technical assistance and to Drs. Raymond C. Stevens, Radostin Danev, Denise Wootten and Patrick M. Sexton for valuable interactions. This work was partially supported by National Natural Science Foundation of China 81872915 (M.-W.W.), 82073904 (M.-W.W.), 32071203 (L.H.Z), 81922071 (Y.Z.), 81773792 (D.Y.), 81973373 (D.Y.) and 21704064 (Q.Z.); National Science and Technology Major Project of China – Key New Drug Creation and Manufacturing Program 2018ZX09735–001 (M.-W.W.) and 2018ZX09711002–002–005 (D.Y.); the National Key Basic Research Program of China 2018YFA0507000 (M.-W.W.) and 2019YFA0508800 (Y.Z.); Ministry of Science and Technology of China 2018YFA0507002 (H.E.X.); Shanghai Municipal Science and Technology Major Project 2019SHZDZX02 (H.E.X.); the Strategic Priority Research Program of Chinese Academy of Sciences XDB37030103 (H.E.X.); Novo Nordisk-CAS Research Fund grant NNCAS-2017–1-CC (D.Y.); Zhejiang Province Science Fund for Distinguished Young Scholars LR19H310001 (Y.Z.); Fundamental Research Funds for Central Universities 2019XZZX001-01-06 (Y.Z.); Novo Shanghai Science and Technology Development Funds 18ZR1447800 (L.H.Z.) and 18431907100 (M.-W.W.); The Young Innovator Association of CAS 2018325 (L.H.Z.) and SA-SIBS Scholarship Program (L.H.Z. and D.Y.). The cryo-EM data were collected at Cryo-Electron Microscopy Research Center, Shanghai Institute of Materia Medica, Chinese Academy of Sciences.

## Author contributions

F.H.Z., Z.T.C., K.N.H. and C.Z. designed expression constructs, purified the receptor complexes, screened the specimen, prepared the final samples for negative staining, collected cryo-EM data and participated in manuscript preparation. X.Y.Z., A.Y.L. and T.X. conducted map calculation and participated in figure preparation; Q.Q.M., M.W., L.N.C. and L.H.Z. built the models of the complexes and carried out structural analyses; Q.T.Z. conducted MD simulations, comparative structural analysis and figure preparation; A.T.D. and Y.C. performed ligand binding and signaling experiments under the supervision of D.H.Y.; R.L.C. and P.Y.X. participated in method development; Y.Z. and B.W. assisted in structural studies on GLP-1R and GCGR; H.E.X. and M.-W.W. initiated the project; Q.T.Z., L.H.Z., H.E.X. and M.-W.W. supervised the studies, analyzed the data and wrote the manuscript with inputs from all co-authors.

### Competing interests

The authors declare no competing interests.

**Fig. S1.**
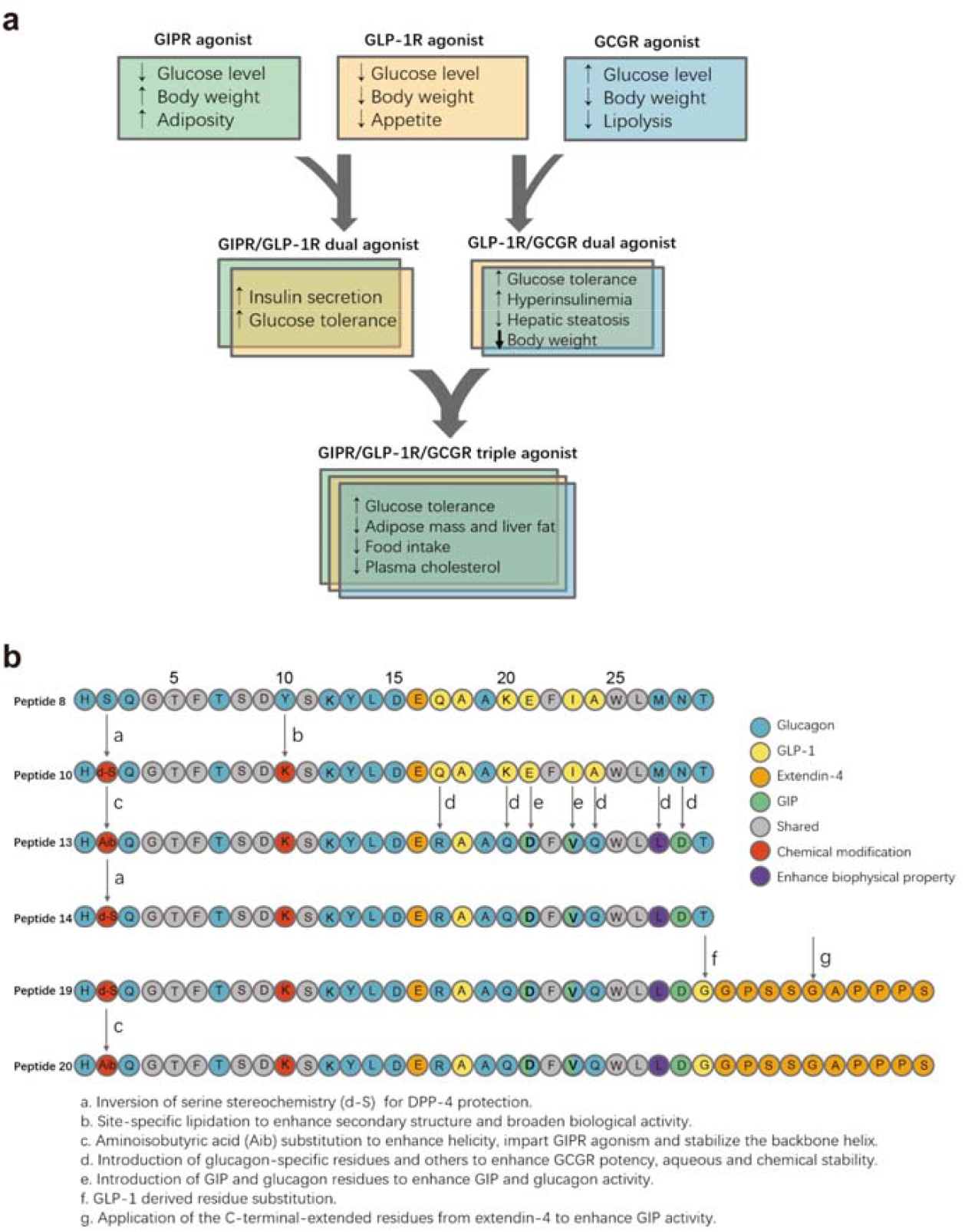
Principles of combinatorial agonism to synergize metabolic actions and maximize therapeutic benefits. **a**, Schematic representation of the therapeutic advantages of dual and triple agonists targeting the human glucose-dependent insulinotropic polypeptide (GIP), glucagon-like peptide-1 (GLP-1) and glucagon (GCG) receptors (GIPR, GLP-1R and GCGR, respectively). GLP-1R agonists are used to treat type 2 diabetes and obesity because of their ability to promote satiety and insulin secretion. Their effect on weight loss could be complemented by that of glucagon on lipolysis and thermogenesis, leading to a series of GLP-1R/GCGR dual agonists (e.g., peptide 8) based on the sequence of GCG. Subsequently, GIPR agonism was added to GLP-1R agonists to enhance the glycemic benefits of GLP-1 resulting in a new series of dual agonists (e.g., tirzepatide) that improved insulin secretion and glucose tolerance while reducing adverse events of the monotherapy. Given the enhanced performance of both dual agonists in the treatment of obesity and T2D, as well as the structural similarity among the three peptides, Unimolecular GIPR/GLP-1R/ GCGR triple agonists (e.g., peptide 20) were developed to combine the strength of both types of dual agonists. **b**, Evolutionary pathway towards a highly potent and balanced unimolecular triple agonist (peptide 20) for GIPR, GLP-1R and GCGR. The modifications and their actions on combinatorial agonism are explained in the bottom.

**Fig. S2.**
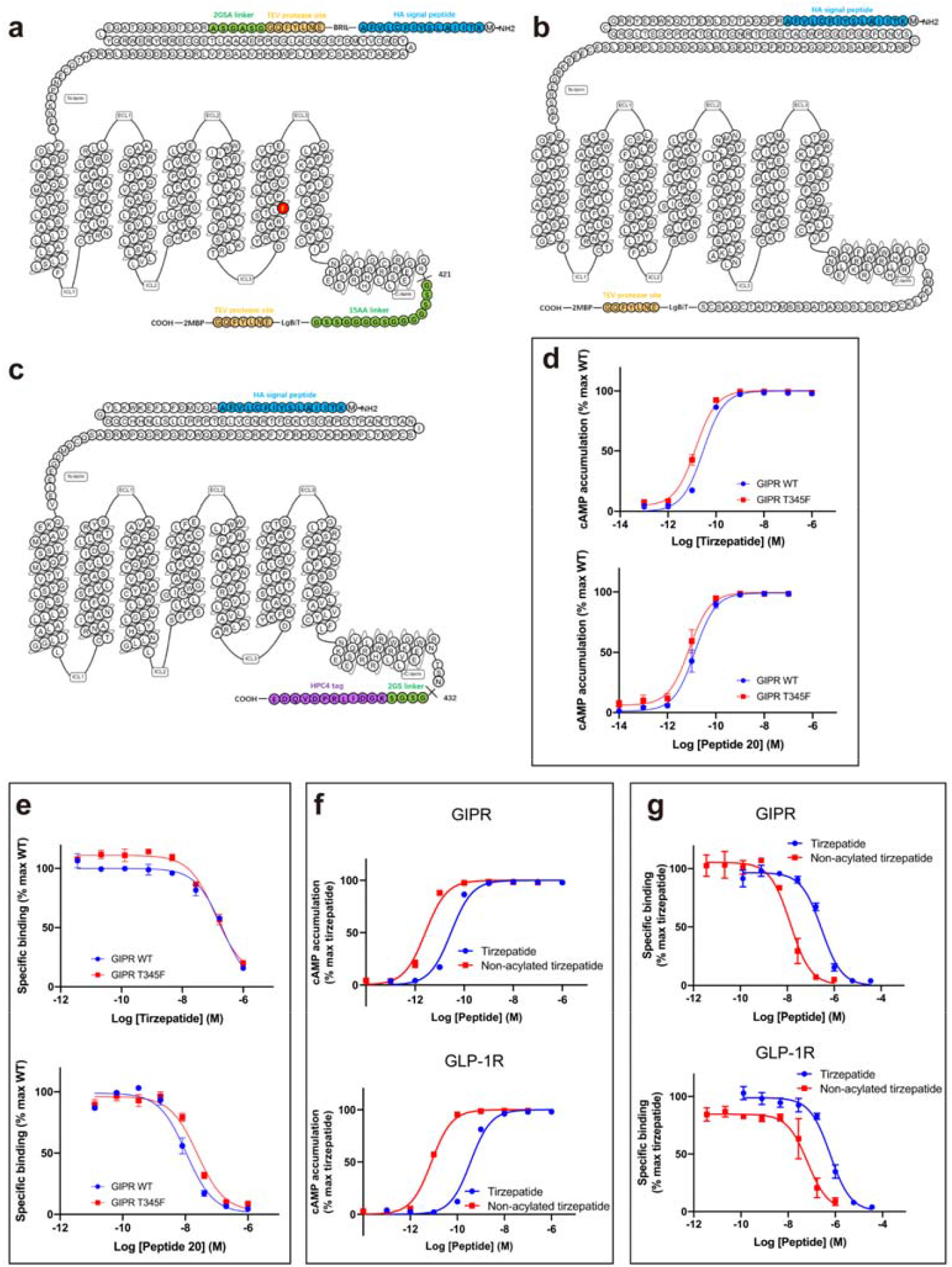
Receptor constructs for structure determination. **a-c**, Schematic diagrams of receptor constructs used for structure determination: GIPR construct (**a**), GLP-1R construct (**b**) and GCGR construct (**c**). **d**, Effects of GIPR T345F on tirzepatide (top) and peptide 20 (bottom)-induced cAMP accumulation. **e**, Effects of GIPR T345F on receptor binding affinities of tirzepatide (top) and peptide 20 (bottom). **f**, Effects of tirzepatide acylation on GIPR (top) and GLP-1R (bottom)-mediated cAMP accumulation. **g**, Effects of tirzepatide acylation on receptor binding affinities with GIPR (top) and GLP-1R (bottom). cAMP accumulation and binding data were normalized to the maximum response of wild-type (WT) or tirzepatide and concentration–response curves were analyzed using a three-parameter logistic equation. The experiments were carried out independently at least twice with similar results.

**Fig. S3.**
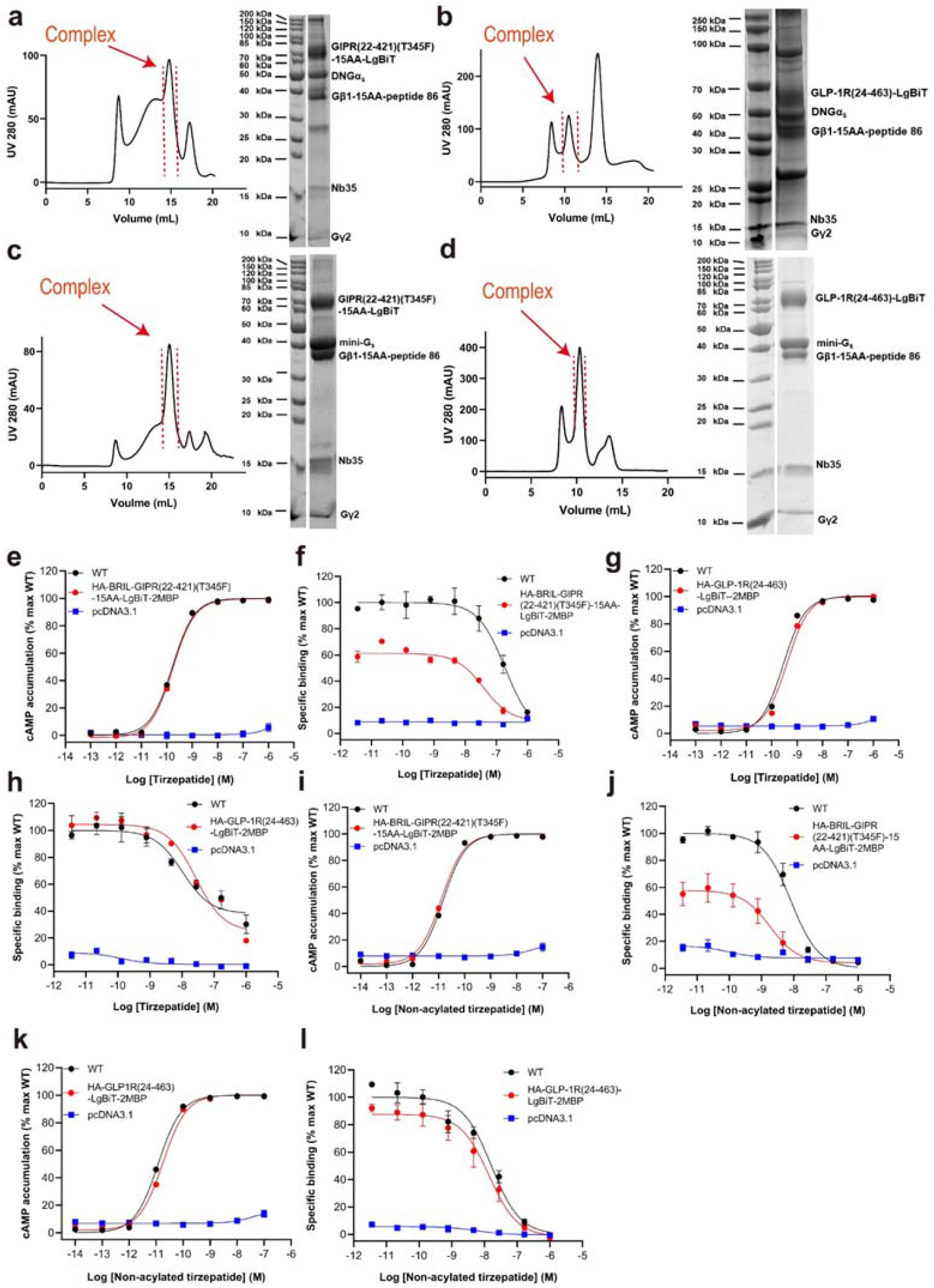
Purification and characterization of the tirzepatide–GIPR/GLP-1R–G_s_–Nb35 complexes and non-acylated tirzepatide–GIPR/GLP-1R–G_s_–Nb35 complexes. **a**, Size-exclusion chromatography on Superose 6 Increase 10/300GL and SDS-PAGE of the tirzepatide–GIPR–G_s_–Nb35 complex. **b**, Size-exclusion chromatography on Superdex 200 Increase 10/300GL and SDS-PAGE of the tirzepatide–GLP-1R–G_s_–Nb35 complex. **c**, Size-exclusion chromatography on Superose 6 Increase 10/300GL and SDS-PAGE of the non-acylated tirzepatide–GIPR-mini–G_s_–Nb35 complex. **d**, Size-exclusion chromatography on Superdex 200 Increase 10/300GL and SDS-PAGE of the non-acylated tirzepatide–GLP-1R–mini-G_s_–Nb35 complex. **e**, cAMP responses following tirzepatide stimulation in HEK 293T cells transfected with wild-type (WT) or modified GIPR constructs. **f**, Binding of tirzepatide to the full-length or modified GIPR in competition with ^125^I-GIP_1-42_. **g**, cAMP responses following tirzepatide stimulation in HEK 293T cells transfected with WT or modified GLP-1R constructs. **h**, Binding of tirzepatide to the full-length or modified GLP-1R in competition with ^125^I-GLP-1_(7-36)_NH_2_. **i**, cAMP responses following non-acylated tirzepatide stimulation in HEK 293T cells transfected with WT or modified GIPR constructs. **j**, Binding of non-acylated tirzepatide to the full-length or modified GIPR in competition with ^125^I-GIP_1-42_. **k**, cAMP responses following non-acylated tirzepatide stimulation in HEK 293T cells transfected with WT or modified GLP-1R constructs. **l**, Binding of non-acylated tirzepatide to the full-length or modified GLP-1R in competition with ^125^I-GLP-1_(7-36)_NH_2_. Signals were normalized to the maximum response of the WT and dose-response curves were analyzed using a three-parameter logistic equation. Whole cell binding assay was performed in CHO-K1 cells. Binding data were analyzed using a three-parameter logistic equation to determine pIC_50_ and span values. Data shown are means ± S.E.M. of at least three independent experiments.

**Fig. S4.**
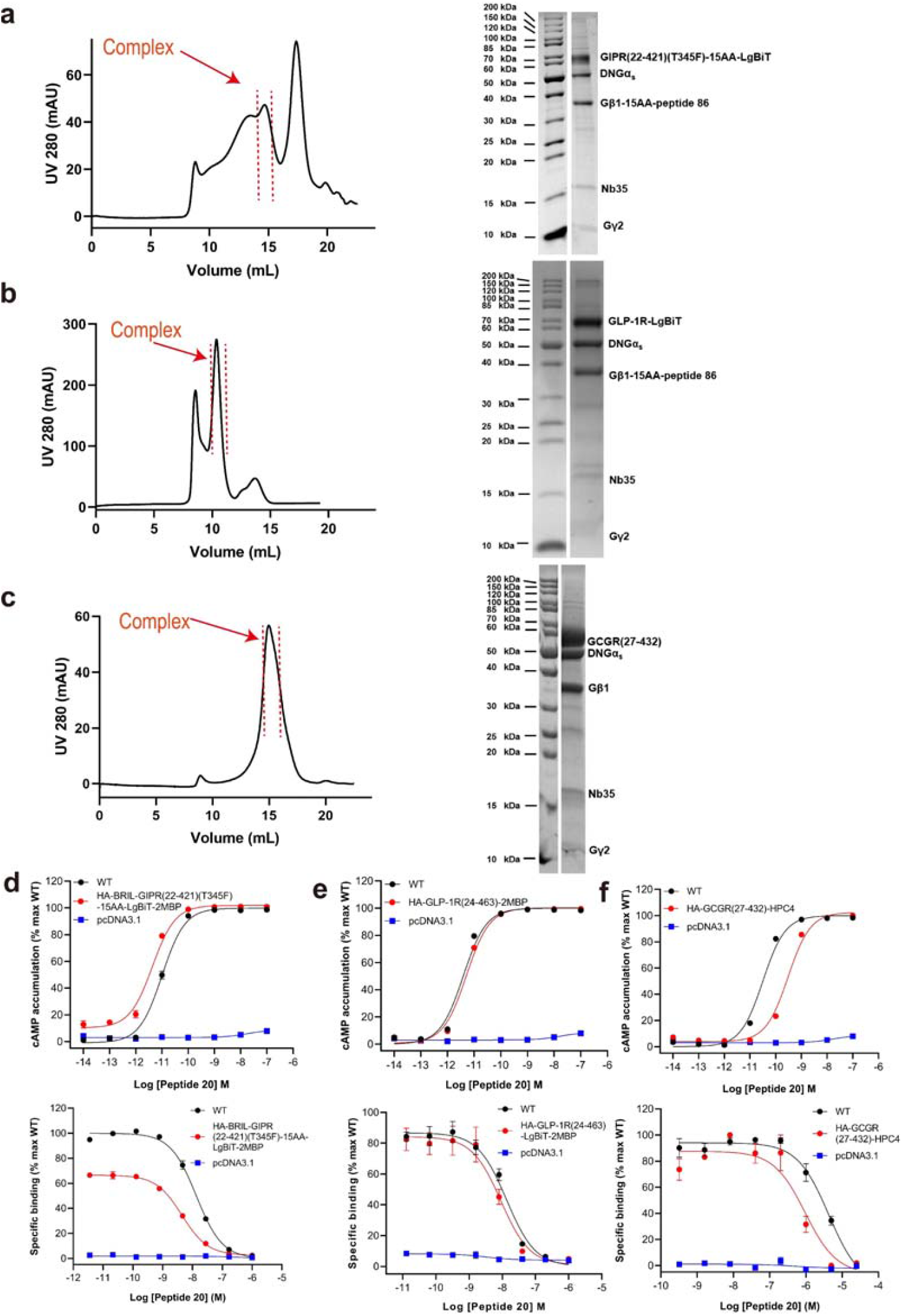
Purification and characterization of the peptide 20–GIPR/GLP-1R/GCGR–G_s_–Nb35 complexes. **a**, Size-exclusion chromatography on Superose 6 Increase 10/300GL and SDS-PAGE of the peptide 20–GIPR–G_s_–Nb35 complex. **b**, Size-exclusion chromatography on Superdex 200 Increase 10/300GL and SDS-PAGE of the peptide 20–GLP-1R–G_s_–Nb35 complex. **c**, Size-exclusion chromatography on Superose 6 Increase 10/300GL and SDS-PAGE of the peptide 20–GCGR–G_s_–Nb35 complex. **d**, Top, cAMP responses following peptide 20 stimulation in HEK 293T cells transfected with wild-type (WT) or modified GIPR constructs. Bottom, binding of peptide 20 to the full-length or modified GIPR in competition with ^125^I-GIP_1-42_. **e**, Top, cAMP responses following peptide 20 stimulation in HEK 293T cells transfected with WT or modified GLP-1R constructs. Bottom, binding of peptide 20 to the full-length or modified GLP-1R in competition with ^125^I-GLP-1_(7-36)_NH_2_. **f**, Top, cAMP responses following peptide 20 stimulation in HEK 293T cells transfected with WT or modified GCGR constructs. Bottom, binding of peptide 20 to the full-length or modified GCGR in competition with ^125^I-GCG. Signals were normalized to the maximum response of the WT and dose-response curves were analyzed using a three-parameter logistic equation. Whole cell binding assay was performed in CHO-K1 cells. Binding data were analyzed using a three-parameter logistic equation to determine pIC_50_ and span values. Data shown are means ± S.E.M. of at least three independent experiments.

**Fig. S5.**
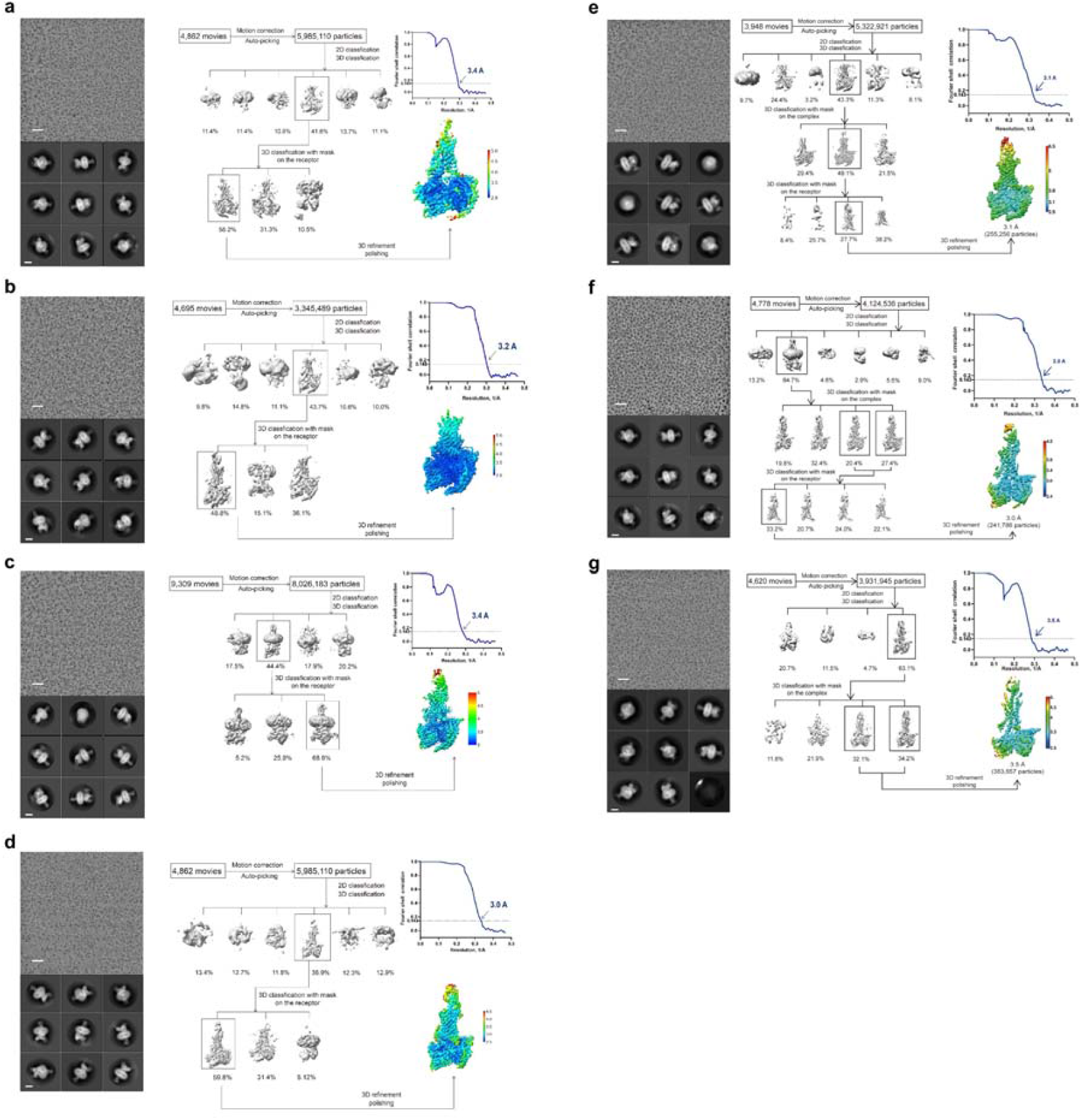
Cryo-EM data processing and validation. **a**, Tirzepatide–GIPR–G_s_ complex: top left, representative cryo-EM micrograph (scale bar: 40 nm) and two-dimensional class averages (scale bar: 5 nm); top right, flow chart of cryo-EM data processing; bottom left, local resolution distribution map of the complex with the ECD and Gold-standard Fourier shell correlation (FSC) curves of overall refined receptor; bottom right, local resolution distribution map of the complex without the ECD and FSC curves of overall refined receptor. **b**, Non-acylated tirzepatide–GIPR–G_s_ complex: left, representative cryo-EM micrograph (scale bar: 40 nm) and two-dimensional class averages (scale bar: 5 nm); middle, flow chart of cryo-EM data processing; right, local resolution distribution map of the complex and FSC curves of overall refined receptor. The experiments were conducted twice independently with similar results. **c**, Tirzepatide–GLP-R–G_s_ complex: left, representative cryo-EM micrograph (scale bar: 40 nm) and two-dimensional class averages (scale bar: 5 nm); middle, flow chart of cryo-EM data processing; right, local resolution distribution map of the complex and FSC curves of overall refined receptor. **d**, Non-acylated tirzepatide–GLP-1R–G_s_ complex: left, representative cryo-EM micrograph (scale bar: 40 nm) and two-dimensional class averages (scale bar: 5 nm); middle, flow chart of cryo-EM data processing; right, local resolution distribution map of the complex and FSC curves of overall refined receptor. The experiments were performed twice independently with similar results. **e**, Peptide 20–GIPR–G_s_ complex: left, representative cryo-EM micrograph (scale bar: 40 nm) and two-dimensional class averages (scale bar: 5 nm); middle, flow chart of cryo-EM data processing; right, local resolution distribution map of the complex and FSC curves of overall refined receptor. The experiments were carried out twice independently with similar results. **f**, Peptide 20–GLP-1R–G_s_ complex: left, representative cryo-EM micrograph (scale bar: 40 nm) and two-dimensional class averages (scale bar: 5 nm); middle, flow chart of cryo-EM data processing; right, local resolution distribution map of the complex and FSC curves of overall refined receptor. The experiments were repeated independently twice with similar results. **g**, Peptide 20–GCGR–G_s_ complex: left, representative cryo-EM micrograph (scale bar: 40 nm) and two-dimensional class averages (scale bar: 5 nm); middle, flow chart of cryo-EM data processing; right, local resolution distribution map of the complex and FSC curves of overall refined receptor. The experiments were executed twice independently with similar results.

**Fig. S6.**
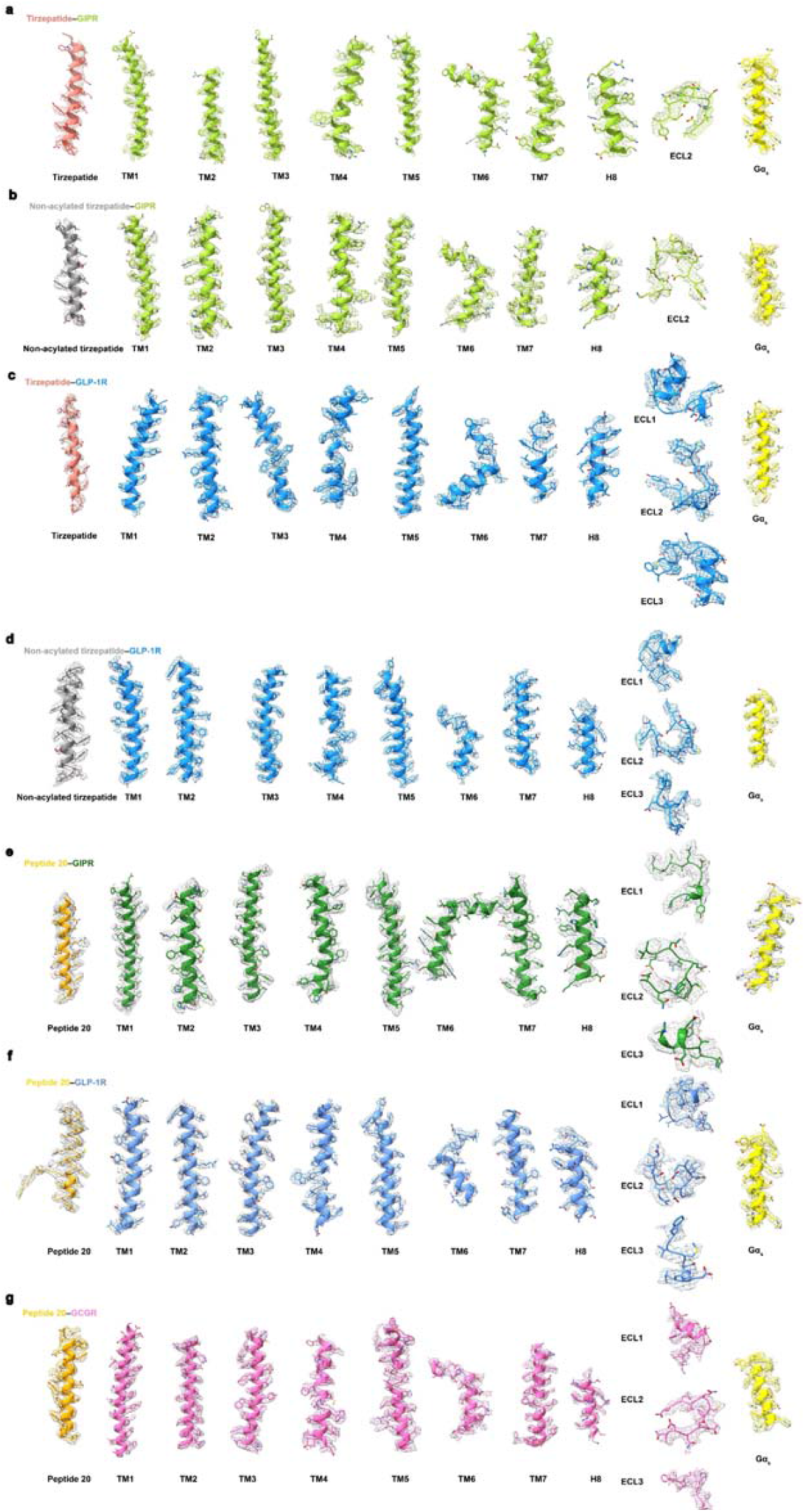
Near-atomic resolution model of the complexes in the cryo-EM density maps. **a**, EM density map and model of the tirzepatide–GIPR–G_s_ complex are shown for all seven-transmembrane α-helices (7TMs), helix 8 and extracellular loop 2 (ECL2) of GIPR, tirzepatide and the α5-helix of the Gα_s_ Ras-like domain. **b**, EM density map and model of the non-acylated tirzepatide–GIPR–G_s_ complex are shown for 7TMs, helix 8 and ECL2 of GIPR, tirzepatide and the α5-helix of the Gα_s_ Ras-like domain. **c**, EM density map and model of the tirzepatide–GLP-1R–G_s_ complex are shown for 7TMs, helix 8 and all extracellular loops of GLP-1R, tirzepatide and the α5-helix of the Gα_s_ Ras-like domain. **d**, EM density map and model of the non-acylated tirzepatide–GLP-1R–G_s_ complex are shown for 7TMs, helix 8 and all extracellular loops of GLP-1R, tirzepatide and the α5-helix of the Gα_s_ Ras-like domain. **e**, EM density map and model of the peptide 20–GIPR–G_s_ complex are shown for 7TMs, helix 8 and all extracellular loops of GIPR, peptide 20 and the α5-helix of the Gα_s_ Ras-like domain. **f**, EM density map and model of the peptide 20–GLP-1R–G_s_ complex are shown for 7TMs, helix 8 and all extracellular loops of GLP-1R, peptide 20 and the α5-helix of the Gα_s_ Ras-like domain. **g**, EM density map and model of the peptide 20–GCGR–G_s_ complex are shown for 7TMs, helix 8 and all extracellular loops of GCGR, peptide 20 and the α5-helix of the Gα_s_ Ras-like domain.

**Fig. S7.**
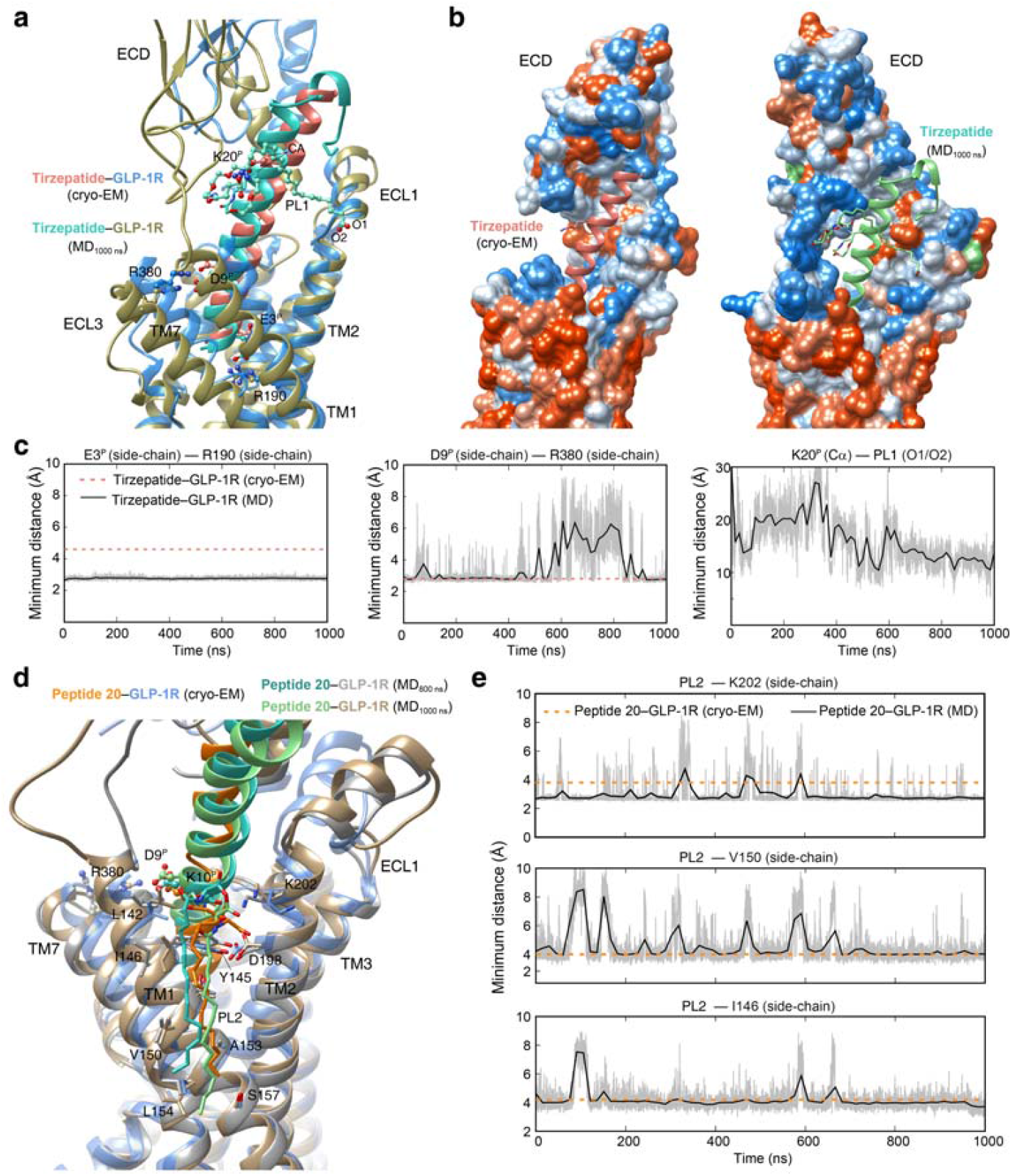
Molecular dynamics (MD) simulation of GLP-1R bound by tirzepatide and peptide 20. **a**, Comparison of tirzepatide conformations between simulation snapshot and the cryo-EM structure. The acylated K20^P^ by a γGlu-2×OEG linker and C18 fatty diacid moiety (named as PL1) is shown in sticks. **b**, Surface representation of the tirzepatide-binding pocket in GLP-1R for cryo-EM structure (left panel) and finial MD snapshot at 1000 ns (right panel). The receptor is shown in surface representation and colored from dodger blue for the most hydrophilic region, to white, to orange red for the most hydrophobic region. **c**, Representative minimum distance between peptide and receptor indicates dynamic conformations of the tail of PL1. **d**, Comparison of peptide 20 conformations between simulation snapshots and the cryo-EM structure. The lipiated K20^P^ by a 16-carbon acyl chain (palmitoyl; 16:0) via a γE spacer (named as PL2), with interacting residues shown in sticks. **e**, Representative minimum distance between heavy atoms of PL2 and its interacting residues suggest that PL2 steadily interacts with the TM1-TM2 crevice residues.

**Fig. S8.**
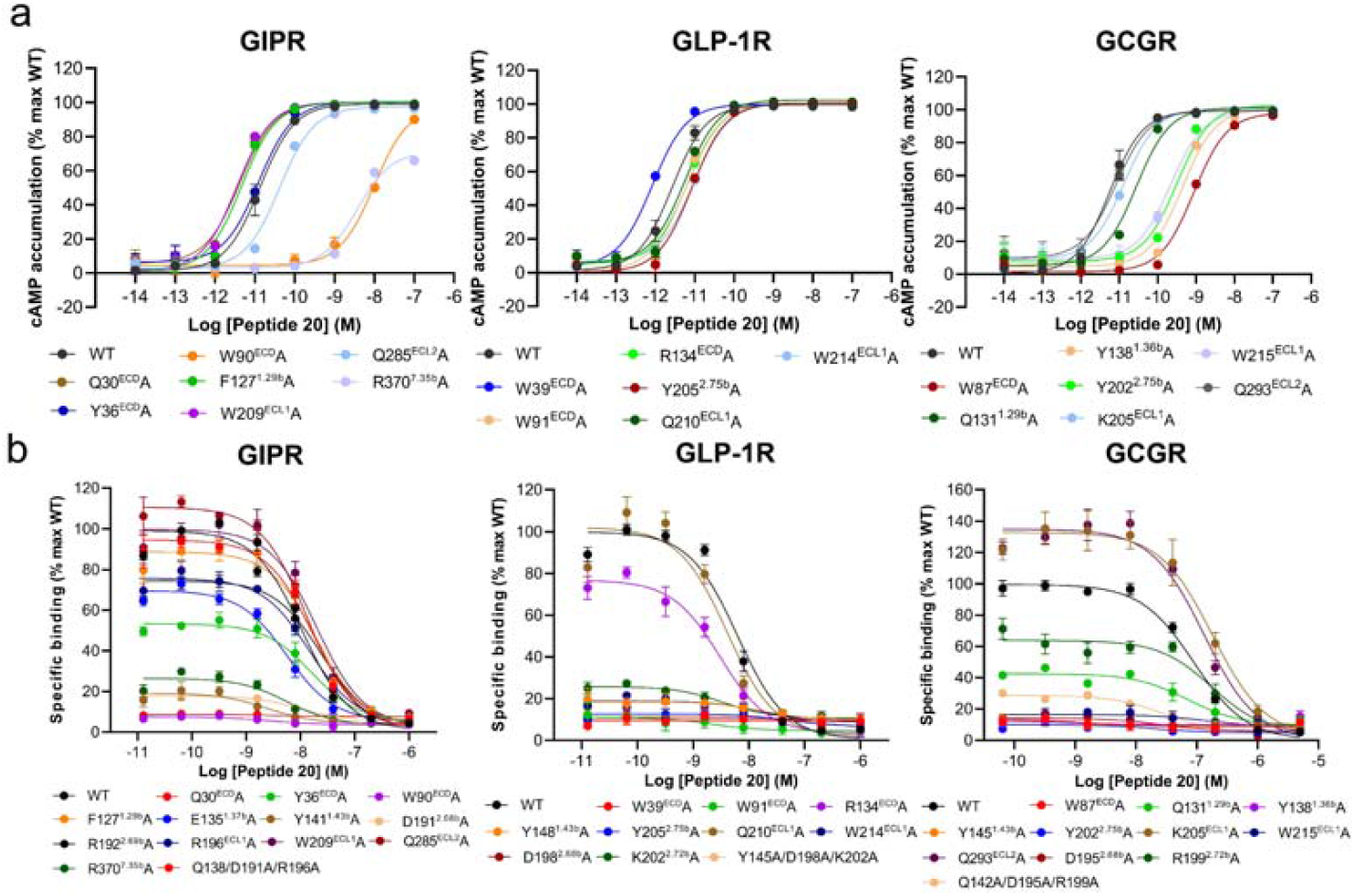
Effect of receptor mutation on peptide 20-induced cAMP accumulation. **a**, Signaling profiles of GIPR (left), GLP-1R (middle) and GCGR (right) mutants. cAMP accumulation was measured in wild-type (WT) and single-point mutated GIPR, GLP-1R or GCGR expressing in HEK 293T cells, respectively. cAMP accumulation was normalized to the maximum response of the WT and dose-response curves were analyzed using a three-parameter logistic equation. Data were generated and graphed as means ± S.E.M. **b**, Binding of peptide 20 to the GIPR (left), GLP-1R (mid) and GCGR (right) mutants in CHO-K1 cells in competition with ^[125I]^-GIP_1-42_, ^125^I-GLP-1_(7-36)_NH_2_ or ^125^I-GCG. Binding data were analyzed using a three-parameter logistic equation to determine pIC_50_ and span values. Data were generated and graphed as means ± S.E.M.

**Figure S9.**
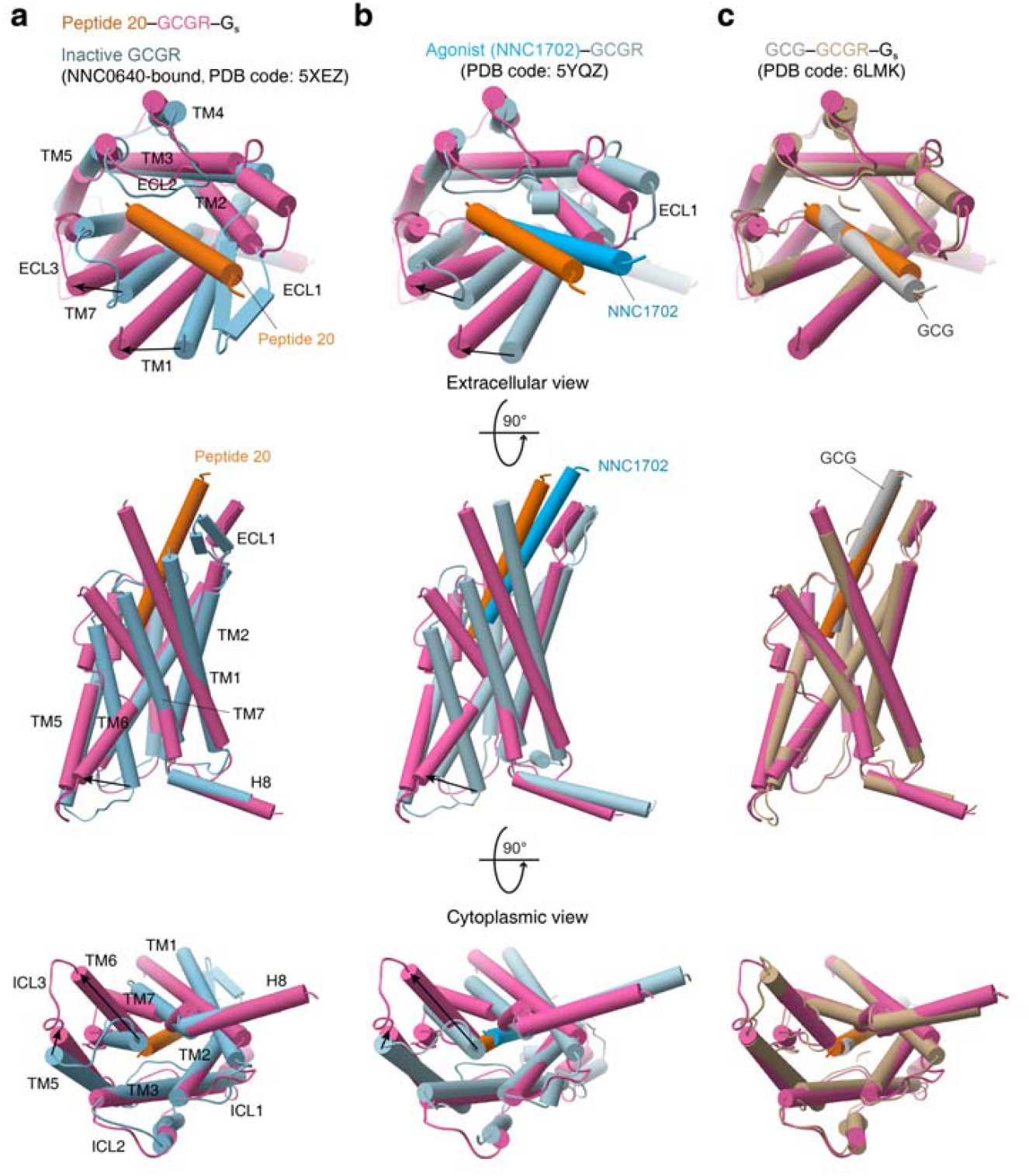
Conformational changes upon GCGR activation. **a-c**, Comparison of peptide 20-bound GCGR with inactive (**a**), agonist-bound (**b**) and both GCG-bound and G protein-coupled active GCGR (**c**). G proteins and receptor ECD are omitted for clarity.

**Table S1.**
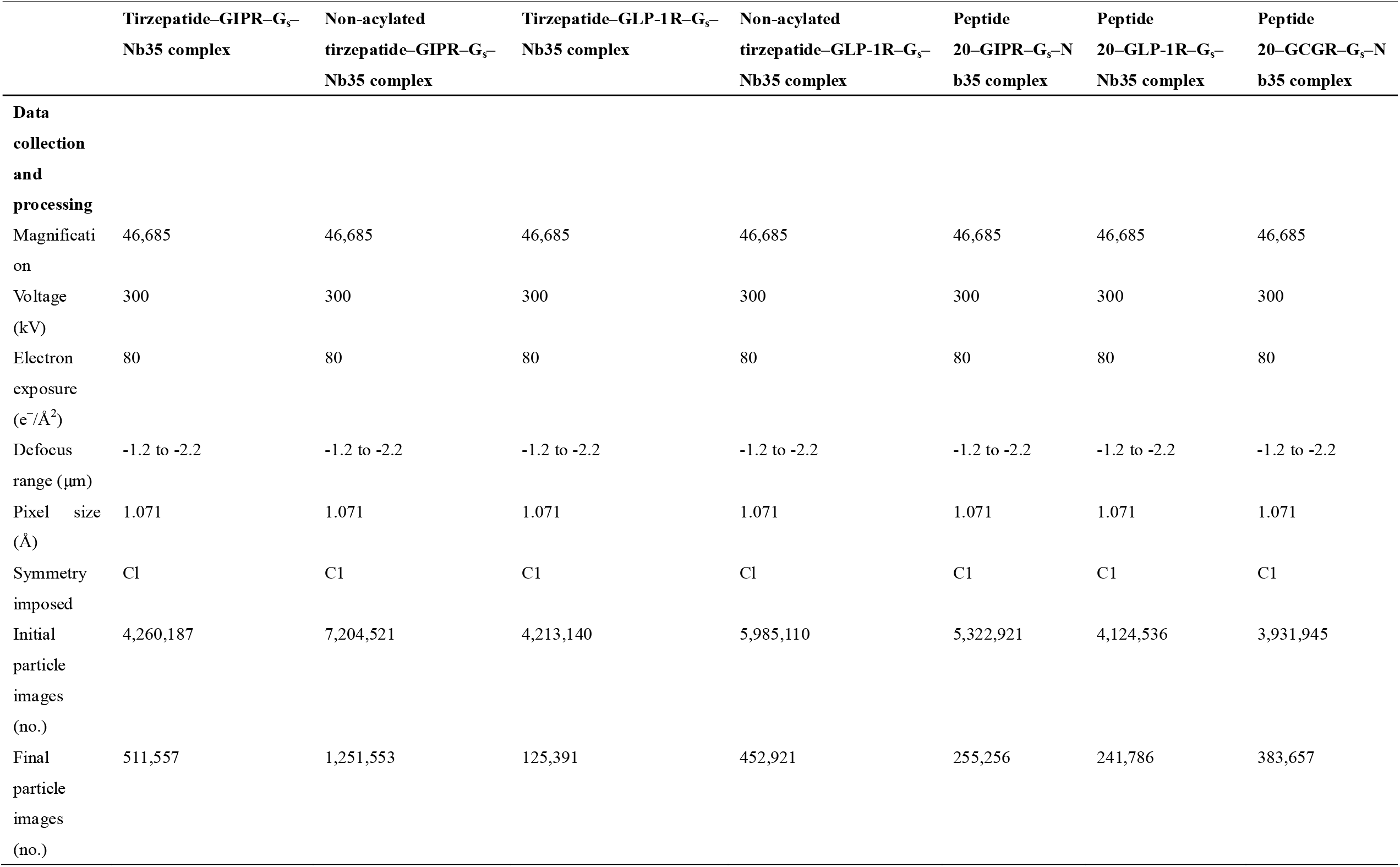
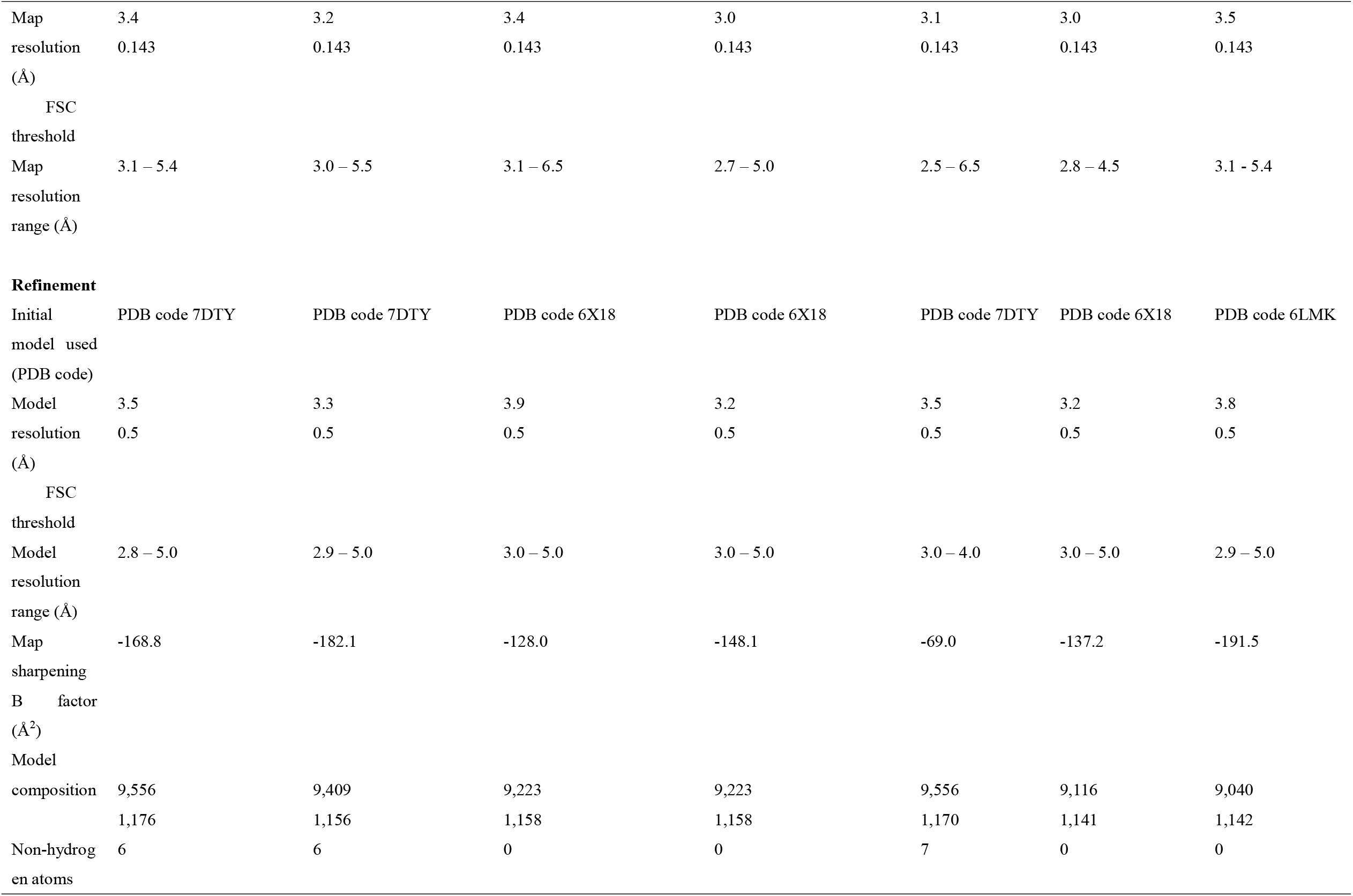
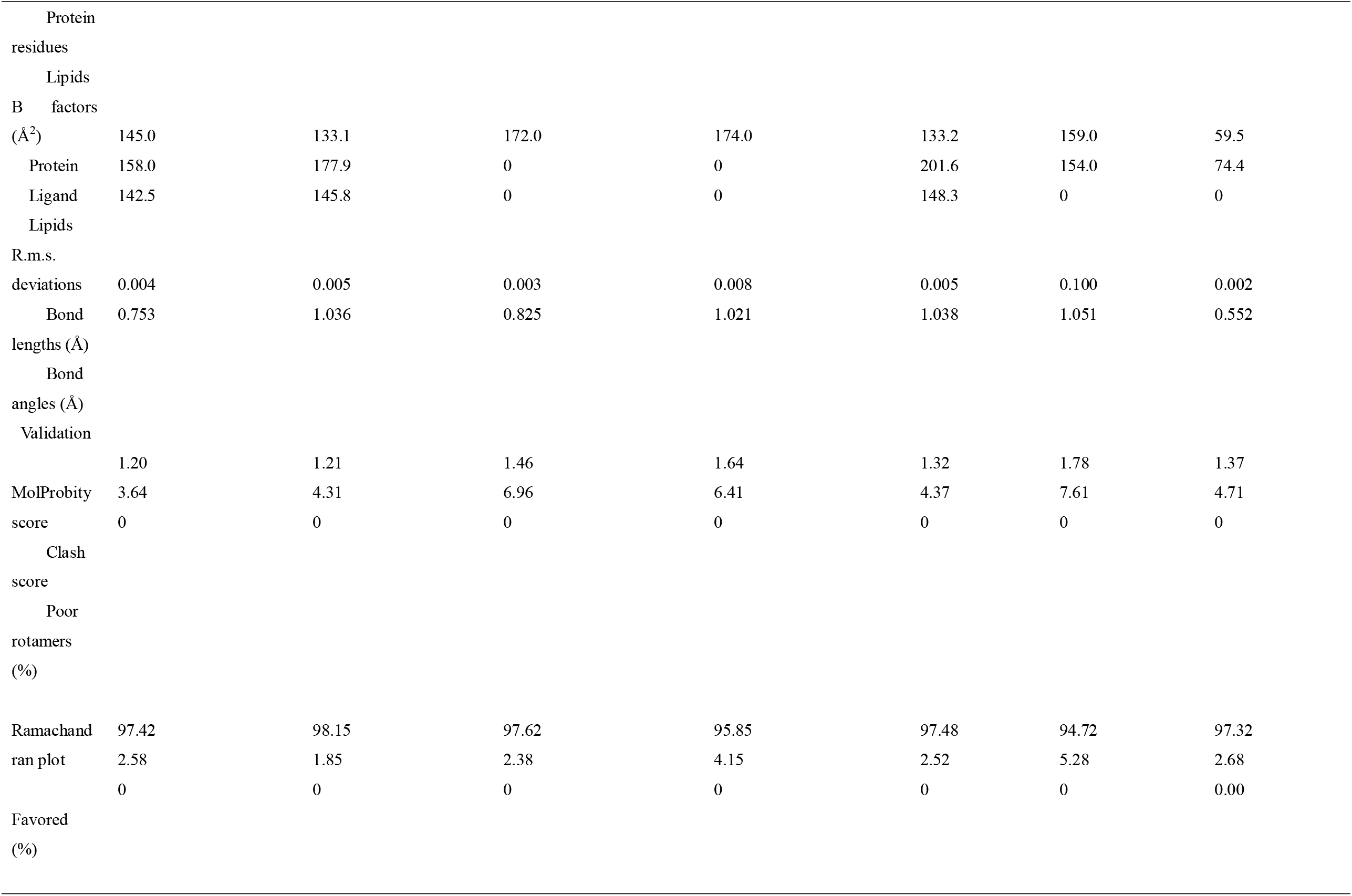
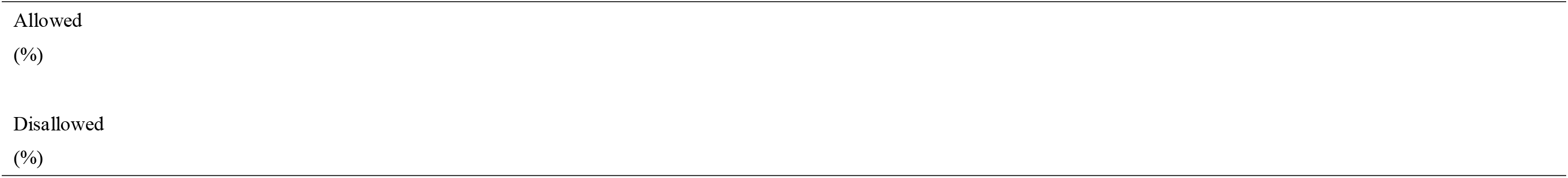
Cryo-EM data collection, refinement and validation statistics.

## Notes

### Competing Interest Statement

The authors have declared no competing interest.

### Summary of Updates

Supplemental files (seven PDBs and their maps) updated

